# *Wolbachia* action in the sperm produces developmentally deferred chromosome segregation defects during the *Drosophila* mid-blastula transition

**DOI:** 10.1101/2022.06.20.496611

**Authors:** Brandt Warecki, Simon Titen, Mohammad Alam, Giovanni Vega, Nassim Lemseffer, Karen Hug, William Sullivan

## Abstract

*Wolbachia*, a vertically transmitted endosymbiont infecting many insects, spreads rapidly through uninfected populations by a mechanism known as Cytoplasmic Incompatibility (CI). In CI, embryos from crosses between *Wolbachia*-infected males and uninfected females fail to develop due to the immediate action of *Wolbachia*-produced factors in the first zygotic division. In contrast, viable progeny are produced when the female parent is infected. Here, we find ∼1/3 of embryos from CI crosses in *Drosophila simulans* develop normally beyond the first and subsequent pre-blastoderm divisions. Developing CI-derived embryos then exhibit chromosome segregation errors during the mid-blastula transition and gastrulation. Single embryo PCR and whole genome sequencing reveal a large percentage of the developed CI-derived embryos bypass the first division defect. Using fluorescence *in situ* hybridization, we find increased chromosome segregation errors in gastrulating CI-derived embryos that had avoided the first division defect. Thus, *Wolbachia* in the sperm induces independent immediate and developmentally deferred defects. Like the initial immediate defect, the delayed defect is rescued through crosses to infected females.

## INTRODUCTION

*Wolbachia* are a bacterial endosymbiont present in the majority of insect species (Weinert et al., 2015; Werren et al., 2008). While they reside in the germline of both sexes, they are vertically transmitted exclusively through the female germline to all the offspring (Kaur et al., 2021). Consequently, *Wolbachia* have evolved a number of strategies that provide a selective advantage to infected females. This includes male killing, conversion of males to fertile females, induction of parthenogenesis, and most commonly cytoplasmic incompatibility (CI) (Serbus et al., 2008). CI is a form of *Wolbachia*-induced conditional sterility. Matings between infected males and uninfected females result in dramatic reductions in egg hatch rates (Hoffmann et al., 1986). However, matings between infected males and infected females, known as the “rescue cross,” results in normal egg hatch rates. Additionally, infected females mated with uninfected males results in normal hatch rates. Thus, in a *Wolbachia*-infected population, infected females have an enormous selective advantage over uninfected females as infected females produce normal hatch rates independent of the infection status of the male (Turelli & Hoffmann, 1991). This phenomenon, as well as *Wolbachia*-induced male sterility, is currently being employed throughout the world as a strategy for combating pest insects and insect-borne human diseases (Jiggins, 2017; Moretti et al., 2018; Zheng et al., 2019).

Since the discovery of CI and rescue (Ghelelovitch, 1952; Yen & Barr, 1971), there have been a number of insights into their molecular and cellular bases (Shropshire et al., 2020). Cytological studies demonstrate failures in condensation, alignment, and segregation of the paternal chromosomes during the first zygotic division in embryos derived from the CI cross (Breeuwer & Werren, 1990; Callaini et al., 1997; Lassy & Karr, 1996; Reed & Werren, 1995; Ryan & Saul, 1968; Tram et al., 2006; Tram & Sullivan, 2002). Subsequent studies demonstrated defects in the protamine-to-histone transition: deposition of the maternally-supplied histone H3.3 is significantly delayed (Landmann et al., 2009). In addition, the male, but not the female, pronucleus of CI-derived embryos exhibits delays in DNA replication, nuclear envelope breakdown, and Cdk1 activation (Tram & Sullivan, 2002). As a result, passage of the male pronucleus through mitosis is delayed relative to the female pronucleus. Molecular insight into the mechanism of CI has come from recent studies demonstrating that a pair of *Wolbachia* genes originating from integrated viral DNA, the CI factors or Cifs, is likely responsible for CI (Beckmann et al., 2017; LePage et al., 2017). One of these genes, *cidB*, encodes a deubiquitylating enzyme and the other, *cinB*, a nuclease (Chen et al., 2020). When the gene pair is expressed in the male germline, paternal chromosome and embryo abnormalities strikingly similar to *Wolbachia*-mediated CI are observed (Beckmann et al., 2017; LePage et al., 2017).

In addition to the well-characterized first division mitotic defects, studies in a number of species demonstrate that CI produces additional developmental defects and lethal phases later in embryogenesis (Bonneau et al., 2018; Callaini et al., 1997; Callaini et al., 1996; Duron & Weill, 2006; Jost, 1970; Lassy & Karr, 1996; Wright & Barr, 1981). Studies in wasps in which fertilized eggs develop into diploid females and unfertilized eggs develop into haploid males, provide insight into the different developmental outcomes in CI crosses (Tram et al., 2006). If CI disrupts but does not prevent paternal chromosome segregation, the resulting aneuploid embryos fail to develop. In contrast, if CI results in the complete failure of paternal chromosome segregation, embryos develop into haploid males bearing only the maternal chromosome complement (Ryan & Saul, 1968; Tram et al., 2006). In diplo-diploid organisms (where haploid embryos do not develop to adulthood), complete failure of paternal chromosome segregation in the first division leads to haploid embryos that develop and then subsequently fail to hatch (Bonneau et al., 2018; Callaini et al., 1997; Callaini et al., 1996; Duron & Weill, 2006; Jost, 1970). Additionally, in some organisms such as *Drosophila simulans*, a small fraction of CI-derived embryos do not undergo any first division defect and consequently hatch as diploids (Lassy & Karr, 1996).

In conjunction with broad developmental abnormalities, embryos developing from CI crosses also experience various cellular defects (Callaini et al., 1996). For example, studies in *Drosophila* have observed chromosome bridging in CI-derived embryos developing through the pre-blastoderm divisions (nuclear cycles 2-9) (Lassy & Karr, 1996; LePage et al., 2017). Additionally, irregular spindles are observed in syncytial and cellularized blastoderms (nuclear cycles 10-14) from CI crosses (Callaini et al., 1996). Other defects in developed embryos from CI-crosses include displaced nuclei, clumped chromatin, and disorganized centrosomes in blastoderms and abnormally condensed nuclei in gastrulating embryos (Callaini et al., 1996). Whether these cellular defects in later stage embryos are the direct result of aneuploidy from first division errors or due to a second, independent set of CI-induced defects is unresolved.

Here, through a combination of live and fixed analysis, we directly address whether late developmental and cellular defects observed in CI-derived embryos are an outcome of the well-characterized first division errors or caused by an independent, poorly understood, set of CI-induced cell cycle defects. Consistent with previous reports, we find that the majority of embryos derived from the CI cross arrest in the first division. About one-third of CI-derived embryos progress normally through the first zygotic and subsequent internal syncytial divisions. We find that ∼40% of the CI-derived embryos that reach the blastoderm stage (>nuclear cycle 10) are diploid, having undergone a normal first division. While these embryos undergo normal pre-blastoderm divisions, they exhibit significantly increased chromosome segregation defects during the mid-blastula transition (MBT), cellularization, and gastrulation. Crosses to infected females (the rescue cross) reduce the frequencies of the induced division errors. Thus, CI produces distinct and independent early and late chromosome segregation defects. These results reveal that *Wolbachia* CI-induced defects in the sperm produce developmentally deferred chromosome segregation defects in the late blastoderm divisions. These findings provide insight into the mechanisms of CI.

## RESULTS

### *Wolbachia*-induced CI produces a late embryonic lethality in addition to early embryonic lethality

We used a combination of fixed and live analyses to determine the timing of defects as CI-derived embryos progressed through the early blastoderm divisions, cellularization, gastrulation, and hatching. We compared four different crosses: 1) the wild-type cross (uninfected male x uninfected female), 2) the CI-inducing cross (infected male x uninfected female), 3) the rescue cross (infected male x infected female), and 4) the reciprocal cross (uninfected male x infected female) (Figure 1A). Unless otherwise noted, we performed all experiments with *D. simulans* stocks that shared the same genetic background and were infected or uninfected with *Wolbachia* (*wRiv*) (see Materials and Methods). We used *D. simulans* because CI is particularly pronounced in this species.

**Figure 1.**
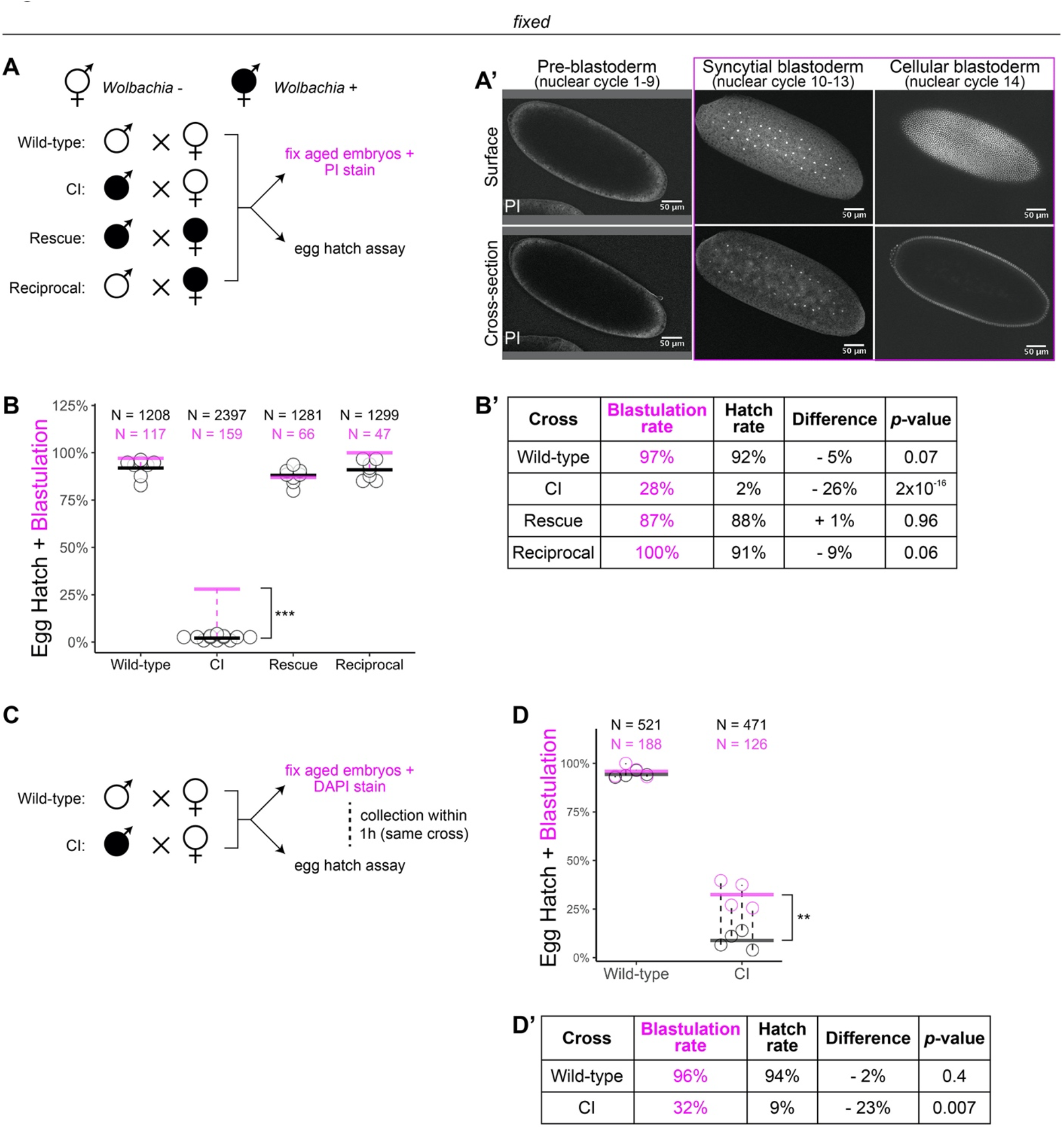
*Wolbachia* induces both early and late embryonic lethality. (A) *Wolbachia* infection status is indicated by filled circles. Embryos were collected from each of the four crosses and either used for egg hatch assays or aged prior to fixing and staining DNA with propidium iodide (PI). (A’) Confocal imaging of PI-stained embryos allowed categorization of embryo stage as pre-blastoderm (cycles 2-9), syncytial blastoderm (cycles 10-13), or cellular blastoderms (cycle 14). Scale bars are 50 µm (B-B’) Comparison between blastulation rate (% of fixed embryos staged as progressing beyond cycle 9) and egg hatch rate between each of the four crosses. Each circle represents one egg hatch assay. Black and magenta lines represent the average egg hatch rate and the blastulation rate respectively. While the hatch rate from wild-type, rescue, and reciprocal crosses closely corresponded to the blastulation rate, the hatch rate from CI embryos was statistically significantly decreased compared to the blastulation rate. (C) Embryos were collected from wild-type and CI crosses and were used to either determine embryo stage (by DAPI staining) or egg hatch percentage in paired assays (collections were from the same crosses within 1h of each other). (D-D’) Comparison between blastulation rate (% of fixed embryos staged as progressing beyond cycle 9) and egg hatch for each cross. Each circle represents an experimental replicate. Dashed lines connect paired experiments. Black and magenta lines represent the average egg hatch rate and the blastulation rate respectively. The difference between blastulation rate and hatch rate was statistically significant by a two-sided paired t-test (D’). See also Figure S1.

To determine the timing of CI-induced embryonic lethal phases, we collected embryos from all four crosses and compared egg hatch rates to the percentage of embryos that had developed to at least nuclear cycle 10 (the syncytial blastoderm stage) (Figure 1A-A’). Consistent with previous results (Hoffmann et al., 1986), we observed a severe decrease in hatching for embryos derived from CI crosses (CI=2%; N=2397, compare to wild-type=92%, N=1208; rescue=88%, N=1281; reciprocal=91%, N=1299) (Figure 1B-B’). Thus, both CI-induced embryonic lethality and its corresponding rescue by maternally supplied *Wolbachia* are robust in *D. simulans*. Our analysis of fixed embryos revealed that the percentage of embryos that had developed to nuclear cycle 10 derived from wild-type (97%, N=117), rescue (87%, N=66), and reciprocal (100%, N=47) crosses matched their respective hatch rates. However, unique to the CI cross, the percentage of CI-derived embryos that had developed to nuclear cycle 10 (28%, N=159) was significantly higher than its hatch rate (2%, *p*=2×10^-16^ by χ-square test) (Figure 1B-B’). Therefore, a second lethal phase occurs at or after cortical nuclear cycle 10 that results in a significant proportion of the reduced egg hatch in CI-derived embryos.

To reduce any biological and environmental factors that could influence CI strength and embryonic development, we collected embryos from the same wild-type and CI crosses within 1h of each other (Figure 1C). We then analyzed the egg hatch rate with one set of embryos while fixing and DAPI-staining the other set to determine developmental stage. As before, the percentages of embryos that had developed to nuclear cycle 10 (96%, N=188) and hatched (94%, N=521) were similar in wild-type crosses (*p*=0.4 by two-sided paired t-test) (Figure 1D-D’). In contrast, in CI crosses, the percentages of embryos that had developed to nuclear cycle 10 (32%, N=126) were significantly higher than the percentages of embryos that hatched (9%, N=471) (*p*=0.007 by two-sided paired-t test) (Figure 1D-D’).

As an independent means of determining the lethal phases of embryos derived from the CI cross, we performed live analysis to compare the proportion of pre-blastoderm (nuclear cycles 2-9), syncytial blastoderm (nuclear cycles 10-13), or cellular blastoderm (nuclear cycles 14) in CI and wild-type crosses (Figure S1A-A’). Wild-type embryos developed to the syncytial blastoderm stage 91% (N=40) of the time and hatched at a rate of 92% (N=58). However, CI-derived embryos developed to the syncytial blastoderm stage 38% (N=147) of the time but hatched at a significantly reduced rate of 16% (N=110, *p*=2×10^-4^ by χ-square test) (Figure S1B-B’).

Therefore, consistent with previous results (Bonneau et al., 2018; Callaini et al., 1997; Callaini et al., 1996; Duron & Weill, 2006), these data suggest at least two distinct lethal stages are associated with CI: the well-described lethal phase immediately following fertilization (∼70% of embryos), and a second lethal phase that occurs well after the nuclei have undergone many rounds of syncytial and cellular mitoses (∼30% of embryos). Significantly, rescue acts on both phases.

### Late-stage CI embryos initially develop normally through pre-blastoderm syncytial divisions before exhibiting defects during blastulation

CI induces defective paternal chromosome segregation during the first embryonic division, which can result in either complete or partial loss of paternal chromosomes (Bonneau et al., 2018; Landmann et al., 2009; Tram et al., 2006). One of the possible consequences of improper paternal chromosome segregation in the first division is daughter nuclei that inherit only part of the paternal chromosomes. This resulting segmental aneuploidy may then carry over into the subsequent mitoses (Lassy & Karr, 1996; LePage et al., 2017). Certainly, persistent DNA damage carried by the paternal chromatin could affect repeated syncytial divisions in the form of breakage-fusion-bridge cycles (McClintock, 1941; Titen & Golic, 2008).

Therefore, to assess any contribution of the first division segregation errors to the late-stage CI-induced lethality, we examined fixed and DAPI-stained embryos in all stages of early embryonic development (nuclear cycle 2-14) (Figure 2A). For embryos fixed during nuclear cycles 2-9, we scored for anaphase bridging, unequally-sized telophase daughter nuclei, and disorganized distributions of syncytial nuclei. For embryos fixed in cycles 10-14, we additionally scored for nuclear fallout, a process in which the products of defective divisions recede below the normal cortical monolayer of nuclei (Sullivan et al., 1990). As expected, wild-type-derived embryos exhibited abnormalities in 0% (0/64) of syncytial pre-blastoderm divisions (cycles 2-9), 0% (0/13) of early cortical divisions (cycles 10-11), and 2% (1/58) of late cortical divisions (cycles 12-14) (Figure 2B). Similarly, CI-derived embryos exhibited abnormalities in only 3% (2/63) of syncytial pre-blastoderm divisions (cycles 2-9). However, we observed a significant increase in CI-derived embryos with abnormal nuclei during early cortical divisions (cycles 10-11, 24%, 26/108) and late cortical divisions (cycles 12-14, 38%, 72/190) (*p*=1.1×10^-5^ by two-sided Fisher’s exact test) (Figure 2B). We regularly observed nuclear fallout (55%; Figure 2A, yellow arrows), anaphase bridging (33%), and disorganized nuclei (11%) in CI-derived cycle 10-14 embryos (Figure 2C). Additionally, we found that the CI-induced increase in abnormal nuclear divisions during late cortical divisions (cycles 12-14) was dramatically, but not completely, reduced in the rescue cross (5%, 7/128) (*p*=2.8×10^-11^ by two-sided Fisher’s exact test) (Figure 2B).

**Figure 2.**
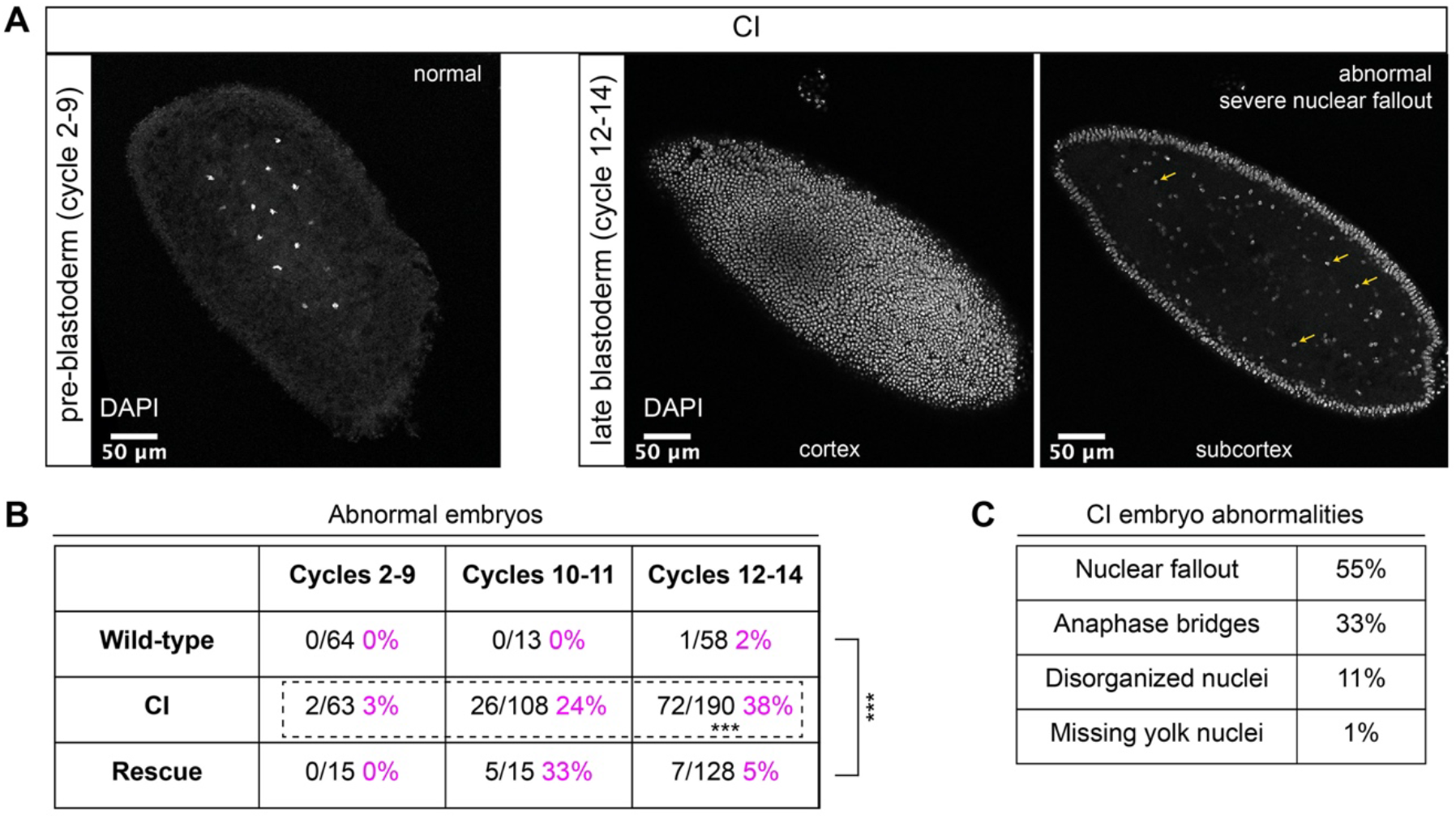
CI-derived embryos proceed normally through pre-blastoderm divisions and then exhibit cellular defects during blastoderm divisions. (A) Examples of fixed and DAPI-stained CI-derived embryos from pre-blastoderm (cycles 2-9), and late blastoderm (12-14) stages. While the CI-derived pre-blastoderm appears normal, the late blastoderm exhibits severe nuclear fallout (nuclei receded from the cortex and into subcortical regions). Arrows point to several examples of fallen out nuclei. Scale bars are 50 µm. (B) Comparison of the percentage of abnormal embryos from wild-type, CI, and rescue crosses during different stages of embryogenesis. While CI-derived embryos developed normally through cycles 2-9, they exhibited significantly increased abnormalities during cycles 10-14. Abnormalities in cycles 10-14 were significantly reduced in embryos from rescue crosses. (C) Classification of abnormalities observed in CI-derived embryos.

Thus, CI-derived embryos that bypass the first lethal phase develop normally through nuclear cycles 2-9 and then exhibit a dramatic increase in abnormal segregation and nuclear organization during the cortical nuclear cycles (10-14). The normal development through nuclear cycles 2-9 suggests that the cortical division defects observed in the CI-derived embryos are not a direct consequence of abnormal first divisions but may instead be separate CI-induced defects. Significantly, as with the first division CI-induced defects, CI-induced cortical defects are rescued when infected males are crossed to infected females.

### Blastoderm embryos from CI crosses have higher rates of nuclear fallout than embryos from wild-type or rescue crosses

To further explore the defects that arise in blastoderm CI embryos, we performed a more sensitive assay to score the number of abnormal cortical nuclei that recede into the interior of the embryo, known as nuclear fallout. Because the fidelity of cortical nuclear divisions is maintained by a mechanism that eliminates the products of abnormal divisions from the cortex (Sullivan et al., 1990), assaying nuclear fallout provides a quantitative measure of cortical division errors (Sullivan et al., 1993). This assay is sensitive due to the lack of gap phases in cortical divisions. Even in wild-type embryos, nuclear fallout is observed at a low level (Sullivan et al., 1993). Consequently, we used this assay to determine the effect of *Wolbachia*-induced CI on the cortical syncytial divisions.

We regularly observed CI embryos with increased numbers of nuclei that had fallen from the cortical layer of nuclei into the subcortex (Figure 3A, magenta arrows). We quantified the amount of nuclear fallout per cycle 10-14 embryo from wild-type (1.3 +/- 2.4, N=85), CI (6.6 +/- 6.4, N=34), rescue (1.7 +/- 3.8, N=60), and reciprocal (0.9 +/- 1.4, N=35) crosses (Figure 3B-B’). The amount of nuclear fallout per embryo was significantly increased in CI embryos compared to wild-type embryos (*p*=4.5×10^-8^ by Mann-Whitney test) and significantly reduced compared to rescue embryos (*p*=1.7×10^-6^ by Mann-Whitney test). Given the increased nuclear density during the final blastoderm cycle, we observed a more pronounced increase in nuclear fallout in cycle 14 CI embryos (CI=11.7, N=14; wild type=1.0, N=64) (Figure 3B’).

**Figure 3.**
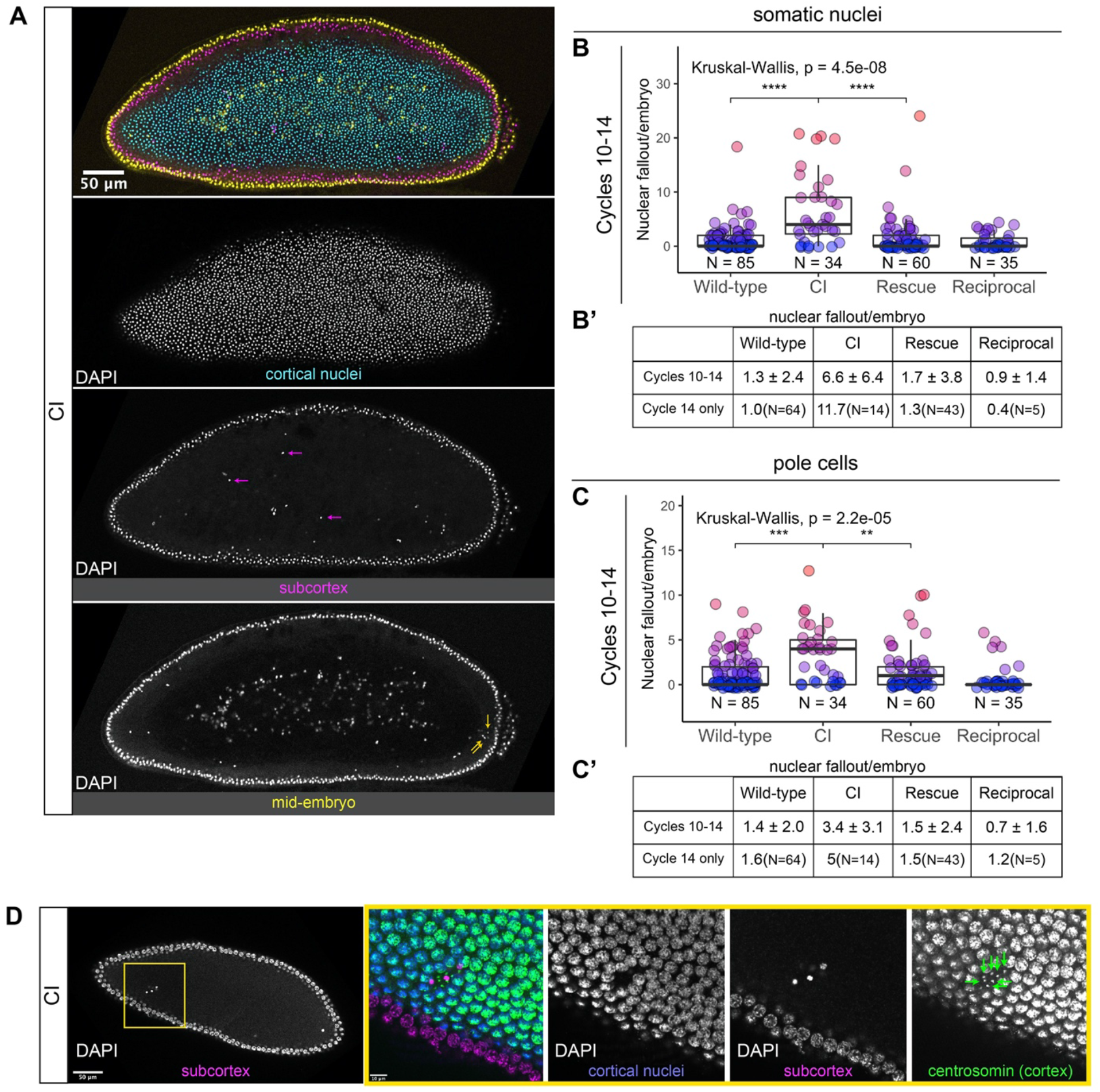
Developed CI-derived embryos exhibit increased rates of nuclear fallout. (A) Image of a CI-derived blastoderm exhibiting moderate nuclear fallout. Cortical nuclei (cyan) are on the surface of the embryo. Nuclei that have fallen out of the cortex can clearly be observed 5-10 µm beneath the cortex (subcortex, magenta) and at the mid-plane of the embryo (yellow). Magenta arrows point to examples of somatic nuclei that have fallen out. Yellow arrows point to examples of pole cells that have fallen out. Scale bar is 50 µm (B-B’) Comparison of somatic nuclear fallout in cycle 10-14 embryos from wild-type, CI, rescue, and reciprocal crosses. (B) Each dot represents the number of fallen nuclei per embryo. (B’) Averages and standard deviations are summarized. CI-derived embryos have significantly increased somatic nuclear fallout compared to wild-type- and rescue-derived embryos. (C-C’) Comparison of pole cell nuclear fallout in cycle 10-14 embryos from wild-type, CI, rescue, and reciprocal crosses. (C) Each dot represents the number of fallen nuclei per embryo. (C’) Averages and standard deviations are summarized. CI-derived embryos have significantly increased somatic nuclear fallout compared to wild-type- and rescue-derived embryos. (D) CI-derived embryo stained with anti-centrosomin antibody to mark centrosomes and counterstained with DAPI. While cortical nuclei (blue) remain strongly associated with their centrosomes (green), nuclei that recede into the subcortex (magenta) detach from their centrosomes that are left at the cortex (green arrows). Yellow box indicates zoomed region. Scales bars are 50 µm and 10 µm for unzoomed and zoomed regions respectively.

The above analyses excluded the extreme posterior region of the embryo. This is because in wild-type embryos, the extreme posterior region is composed of 8-10 cellularized pole cells that have migrated to the cortex ahead of the main contingent of dividing nuclei. These cells are the precursors to the germline (Illmensee & Mahowald, 1974). In general, cortical nuclei in this posterior region exhibit a higher rate of nuclear fallout compared to somatic nuclei in the rest of the embryo (Figure 3A, yellow arrows). Similar to nuclear fallout in the rest of the embryo, nuclear fallout in the posterior pole was dramatically higher in cycle 10-14 embryos from CI crosses compared to those from wild-type, rescue, and reciprocal crosses (Figure 3C-C’).

Previous work has shown that nuclear fallout occurs via detachment of the cortical nuclei from their centrosomes (Sullivan et al., 1993). To determine if the nuclear fallout in embryos from CI crosses is due to a similar detachment from centrosomes, we next fixed embryos from CI crosses and co-stained with DAPI and antibodies that recognize centrosomin, a key component of centrosomes (Megraw et al., 1999) (Figure 3D). Receding nuclei create a gap in the normally dividing cortical surface nuclei. The centrosomes associated with the fallen nuclei (green arrows) remained on the cortex (Figure 3D). Thus, nuclei in CI embryos regularly detach from their centrosomes and recede from the cortex.

### Lagging chromosomes are a proximal cause of nuclear fallout in CI-derived embryos

To determine the primary cause of the errors leading to nuclei falling out from the cortical monolayer of normally dividing nuclei, we injected early embryos with rhodamine-labeled histones and performed live imaging on a confocal microscope. Live analysis allowed us to identify receding nuclei and analyze the proximal mitotic defects that led to nuclear fallout. For both CI- and wild-type-derived embryos, we observed nuclear fallout occurred almost exclusively during the telophase-to-interphase transition (Figure 4A). This is likely the result of a failure of the nuclei maintain association with their centrosomes following errors in the preceding division.

**Figure 4.**
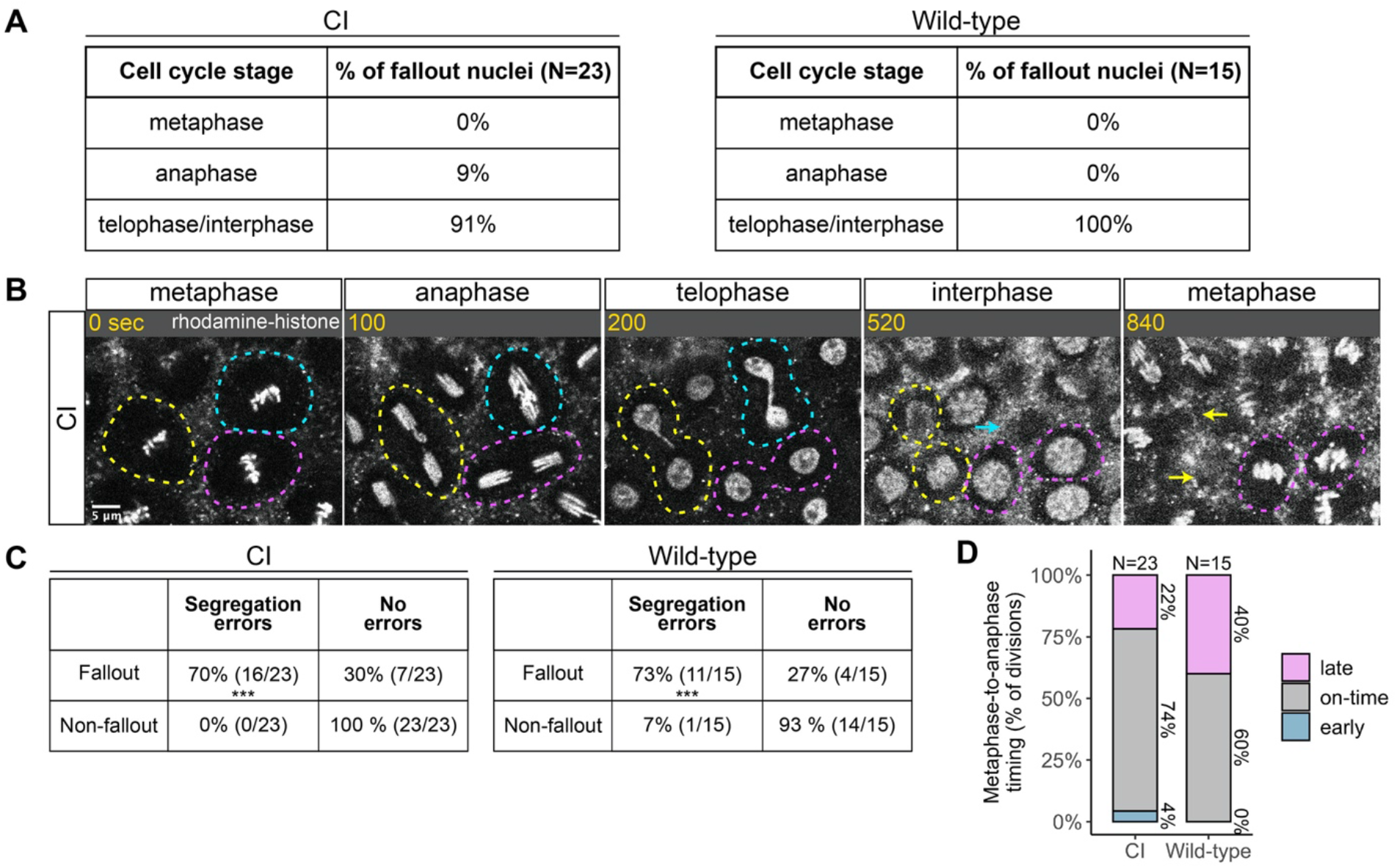
Chromosome segregation errors are the proximate cause of nuclear fallout in CI-derived embryos. (A) Comparison of when in the cell cycle nuclei fallout in both CI- and wild-type-derived embryos. (B) Nuclei that fallout (yellow and blue circles) exhibit severely lagging chromosomes in the previous division, while nuclei that remain at the cortex (magenta circle) exhibit normal chromosome segregation. Scale bar is 5 µm, and time is written in sec. See also Movie 1. (C) Comparison of chromosome segregation errors between nuclei destined to fallout and their neighbors that remain at the cortex (“non-fallout”) in both CI- and wild-type-derived embryos. (D) Comparison of metaphase-to-anaphase timing between nuclei destined to fallout and their neighbors that remain at the cortex in both CI- and wild-type-derived embryos. “Early” = fallout nuclei enter anaphase before their neighbors. “On-time” = fallout and neighboring nuclei enter anaphase simultaneously. “Late” = fallout nuclei enter anaphase after their neighbors. See also Figure S2.

We routinely observed that nuclear fallout in telophase/interphase was immediately preceded by defective sister chromatid separation during anaphase (Figure 4B, Movie 1). While nuclei in which sister chromatids had segregated normally (Figure 4B, magenta circles) remained on the surface and entered the next cell cycle, nuclei in which sister chromatids had severely lagged (Figure 4B, yellow and blue circles) immediately receded into the interior of the embryo during the subsequent interphase. In total, 70% of nuclei destined to fallout in CI-derived embryos were preceded by lagging or bridged chromosomes, as in wild-type-derived embryos (Figure 4C).

Defects causing segregation errors in nuclei destined to fallout may also result in the activation of the spindle assembly checkpoint that would have subsequently delayed entry into anaphase. Therefore, we compared the timing of the metaphase-to-anaphase transition in divisions that resulted in fallout to those of their neighboring normal divisions (Figure 4D). Only a small fraction of nuclei destined to fallout (22%) exhibited a delay in anaphase entry compared to their neighboring nuclei (“late”). In contrast, the vast majority entered anaphase synchronously with (74%, “on-time”) or preceding (4%, “early”) their neighbors. Interestingly, in wild-type embryos, a greater fraction (40%) of nuclei destined to fallout delayed metaphase exit. Therefore, we were unable to regularly detect spindle-assembly checkpoint-mediated delays in CI embryos at our level of temporal resolution.

### Segregation defects persist after cellularization in CI-derived embryos

Following completion of nuclear cycle 13, in an event known as cellularization, each syncytial nucleus is encompassed by an ingressing plasma membrane resulting in the simultaneous formation of individual cells (Sokac et al., 2022). After cellularization, gastrulation begins (Foe, 1989). Invaginations form the head furrow, and bilateral groups of cells throughout the embryo, referred to as mitotic patches, undergo another round of mitosis. We reasoned that CI-induced segregation defects might persist in these mitoses following cellularization.

To examine if chromosome segregation defects occur in CI-derived embryos after the establishment of individual cells, we fixed and DAPI-stained gastrulating embryos from wild-type, CI, and rescue crosses (Figure S2A) and quantified the frequency of division errors in each cross (Figure S2B-D’). While chromosome segregation defects in gastrulating embryos from wild-type crosses occurred at a low rate (11%, N=321 divisions/25 embryos), CI-derived embryos exhibited a significant increase in segregation defects (34%, N=687 divisions/40 embryos) (*p*=7.7×10^-7^ by Mann-Whitney test) (Figure S2D-D’). Significantly, we observed a reduction in segregation errors in embryos from the rescue cross (19%, N=485 divisions/30 embryos) (*p*=8.6×10^-4^ by Mann-Whitney test), although the level of segregation errors was still increased compared to that of wild-type embryos (*p*=0.009 by Mann-Whitney test). Thus, CI-derived embryos exhibit increased chromosome segregation errors that begin in blastoderm stages and continue post-cellularization.

### CI-derived blastoderm embryos consist of both haploids strongly associated with embryonic lethality and diploids

Previous studies have linked late embryonic lethality to haploid development arising from CI-induced chromosome segregation defects during the first division (Bonneau et al., 2018; Callaini et al., 1997; Duron & Weill, 2006). Should CI be strong, paternal chromosomes are completely eliminated during the first division, and embryos develop bearing only the maternal chromosome complement (Tram et al., 2006). In diplo-diplo organisms, these haploid embryos then die before hatching (Bonneau et al., 2018; Callaini et al., 1997; Duron & Weill, 2006). Our observation of mitotic defects in CI-derived blastoderm and gastrulating embryos offers a potential additional explanation for late embryonic lethality. Therefore, we wished to reexamine the relationship between complete paternal chromosome exclusion resulting in haploidy and late embryonic lethality.

To assess the relationship between haploidy and lethality in CI-derived embryos, we performed CI and rescue crosses in which the *Wolbachia*-infected fathers transmitted an *egfp* transgene to all their offspring (Figure 5A). The resulting embryos from these crosses should be genotypically identical. We additionally performed wild-type crosses with uninfected fathers bearing no transgene, serving as a negative control (Figure 5A). We selected embryos that developed to the cellular blastoderm stage and performed single embryo PCR with primers complementary to the paternally-transmitted *egfp* transgene. The *egfp* transgene was always detected in embryos from the positive control rescue cross (∼1.4kb band) and never detected in our negative control embryos derived from uninfected males lacking the *egfp* transgene (Figure 5B). In contrast, we only detected the *egfp* transgene in 42% (N=91) of CI-derived cellular blastoderms (Figure 5B-C). This indicates that while many of the CI-derived blastoderm embryos are diploid, a significant proportion of late-stage CI embryos are haploid. We regularly observed that the overall percentage of *egfp*-positive, diploid embryos correlated with the percentage of hatched eggs from paired egg hatch assays (Figure 5C’), suggesting only the *egfp*-negative, haploid embryos were failing to hatch. Thus, as previously reported (Bonneau et al., 2018; Callaini et al., 1997; Duron & Weill, 2006), haploidy due to loss of paternal chromosomes is linked with late embryonic lethality.

**Figure 5.**
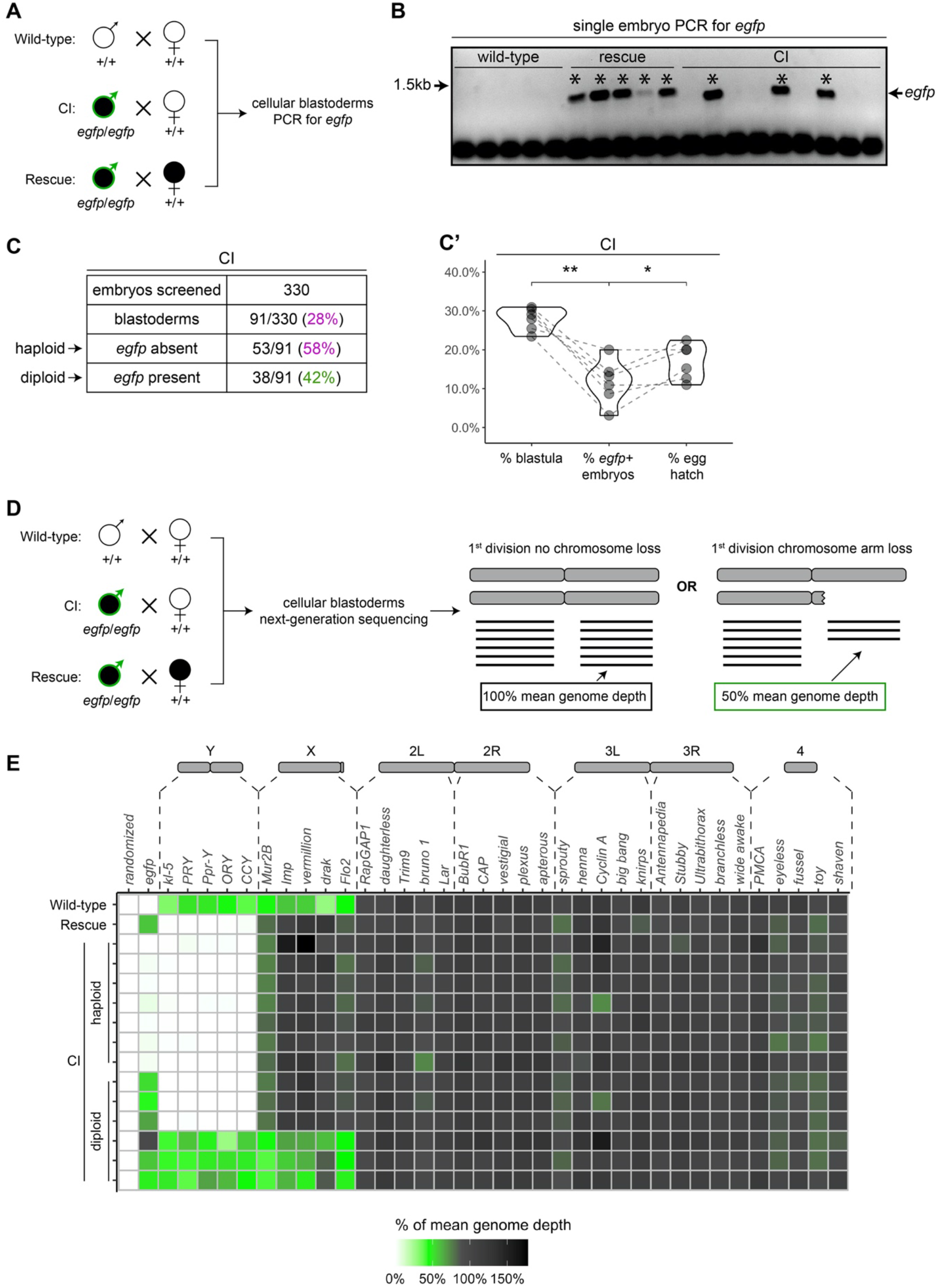
Late-stage CI-derived embryos are either haploid (maternal chromosome set) or diploid (both parental chromosome sets) (A) Embryos were collected from wild-type crosses or CI and rescue crosses in which the father was homozygous for an *egfp* transgene. Embryos were staged live, and cellular blastoderms were selected for single embryo PCR analysis with primers recognizing *egfp*. (B) A representative gel showing detection of *egfp* (asterisks) in all rescue-derived embryos and in a mix of CI-derived embryos. No *egfp* is detected in the wild-type control. (C) Summary of the percentage of screened CI-derived cellular blastoderms in which either *egfp* was absent (haploid) or *egfp* was present (diploid). (C’) Comparison of the percentages of embryos that had reached at least cycle 10 (% blastula), had detectable *egfp* bands (% *egfp*+ embryos) and a concomitant egg hatch (% egg hatch). Each dot represents one experimental replicate, and lines connect values for the same experiment. The percentage of *egfp*+ embryos (diploids) was more associated with the percentage of eggs that hatched (*p*=0.045 by Mann-Whitney test) than with the percentage of blastoderms screened (*p*=0.001 by Mann-Whitney test), suggesting haploid embryos do not hatch. (D) Embryos were collected from wild-type crosses or CI and rescue crosses in which the father was homozygous for an *egfp* transgene. Embryos were staged live, and cellular blastoderms were selected for single embryo sequencing. If chromosome arms were not lost during the first division, the mean depth of reads mapping to the chromosome arm should be near the mean depth of reads mapping to the genome (black box). If chromosome arms were lost during the first division, the mean depth of reads aligning to that chromosome arm should be 50% of the mean depth of reads aligning to the genome (green box). In haploids, maternal chromosome arm loss would result in no reads mapping to that chromosome arm. (E) Sequenced embryos were sorted as haploid or diploid based on the depth of reads mapping to *egfp*. Each box represents the mean depth of reads aligning to that gene divided by the mean depth of reads aligning to the whole genome (“mean genome depth”). White = 0% of mean genome depth; green = 50% of mean genome depth; grey = 100% of mean genome depth; black = 150% of mean genome depth. Consistent with no partial chromosome/chromosome arm loss, genes across all chromosomes were present at 100% mean genome depth for both haploids and diploids (or 50% for X-linked genes when embryos are male). See also Figure S3. See also File S1.

### Late-stage defects are not due to chromosome fragmentation and mosaicism after the first division

Although we found haploidy to be strongly associated with late embryonic lethality, haploidy does not inherently cause the type of chromosome segregation errors we regularly observed in late CI-derived embryos (Debec, 1978; Tang et al., 2017). An alternative potential cause for the segregation errors characterized here is segmental aneuploidy due to an incomplete exclusion of the paternal chromosomes during the first division that does not disrupt early embryonic development. In this scenario, partial chromosome loss or chromosome fragmentation is transmitted from the first division through seemingly normal syncytial divisions and then causes the segregation errors seen in later developmental stages.

To test the possibility that fragmented paternal chromosomes are transmitted through the syncytial divisions, we sequenced the entire genome of collected cellular blastoderms and then mapped read depth to specific coding regions spanning the length of each chromosome (Figure 5D). As above, males in the CI and rescue crosses were homozygous for the *egfp* transgene, allowing us to distinguish between embryos in which paternally-derived chromosomes were either present or absent. We then compared read depth at each locus to the mean read depth across the entire genome for each embryo. These data suggest that neither haploid (*egfp*-negative) nor diploid (*egfp*-positive) CI-derived embryos exhibited whole or partial chromosome loss consistent with the mitotic transmission of only part of the paternal genome (Figure 5E, File S1). Instead, both haploid and diploid embryos had full euploid sets of chromosomes corresponding to either 1n or 2n respectively. This indicates that late-stage CI embryos had either 1) lost all their paternal chromosomes during the first division or 2) did not experience any first division defect at all. Thus, defects observed in late-stage CI embryos cannot be due to partial chromosome loss or fragmentation carried over from the first division.

A separate, potentially interesting, outcome of this experiment is that we found CI-derived embryos regularly had less depth of reads mapping to their entire genome than either wild-type or rescue embryos (Figure S3). This was true for both haploid and diploid CI embryos. Normalizing the depth of reads aligning to the whole genome to the depth of reads aligning to the mitochondrial genome (which should be unchanged) for each embryo suggested differences in sequencing input may not fully explain the decrease in reads mapping to CI embryos (Figure S3B). Although we cannot exclude how any variation in sequencing multiple samples may affect these results, this finding raises the intriguing possibility of intrinsic differences in the chromatin of CI and wild-type-derived blastoderm embryos.

### Late-stage mitotic errors in diploid CI-derived embryos are due to a second CI-induced defect separate from the first division defect

Given neither haploidy nor chromosome fragmentation arising from the first division defect explains the mitotic errors we observed in CI-derived blastoderms and gastrulating embryos, we hypothesized that there is instead a second, CI-induced defect completely separate from the first division defect. To test this hypothesis, we asked if CI embryos that had completely “escaped” the first division defect had increased mitotic errors during later developmental stages.

As described above, late-stage embryos are either haploid, missing their complete paternal chromosome complement, or diploid, having escaped any first division defect to develop with both maternal and paternal chromosome sets (Figure 5E). These diploid embryos can be identified by the presence of a paternally-derived chromosome, such as the Y chromosome, which is only detectable in diploids and never in haploids (Figure 5E). The *D. simulans* Y chromosome can be identified by fluorescence in situ hybridization with Y-specific probes (Ferree & Barbash, 2009). Therefore, we fixed gastrulating embryos from wild-type, CI, and rescue crosses, labelled the Y chromosome with fluorescent probes to select embryos that had escaped the first division defect, counterstained with DAPI to score any mitotic defects.

While Y-bearing gastrulating embryos from wild-type crosses (Figure 6A) exhibited relatively normal chromosome segregation, we observed lagging and bridging chromosomes in Y-bearing embryos from CI crosses (Figure 6B, white arrow). Additionally, in Y-bearing embryos from rescue crosses, chromosome segregation proceeded normally (Figure 6C). The increase in chromosome segregation errors in Y-bearing CI-derived embryos (15%, N=1095 divisions/23 embryos) compared to Y-bearing wild-type embryos (7%, N=1418 divisions/40 embryos) was statistically significant (*p*=1.3×10^-5^ by Mann-Whitney test) (Figure 6D-D’). As the diploid, Y-bearing embryos had completely escaped the first division defects, these results demonstrate that late-stage mitotic errors are due to a second CI-induced defect independent of the first division defect. The reduction of chromosome segregation errors in Y-bearing rescue-derived embryos (8%, N=510 divisions/18 embryos) compared to Y-bearing CI-derived embryos was also statistically significant (*p*=6.6×10^-4^ by Mann-Whitney test) (Figure 6D-D’), indicating maternally supplied *Wolbachia* also rescue this second defect.

**Figure 6.**
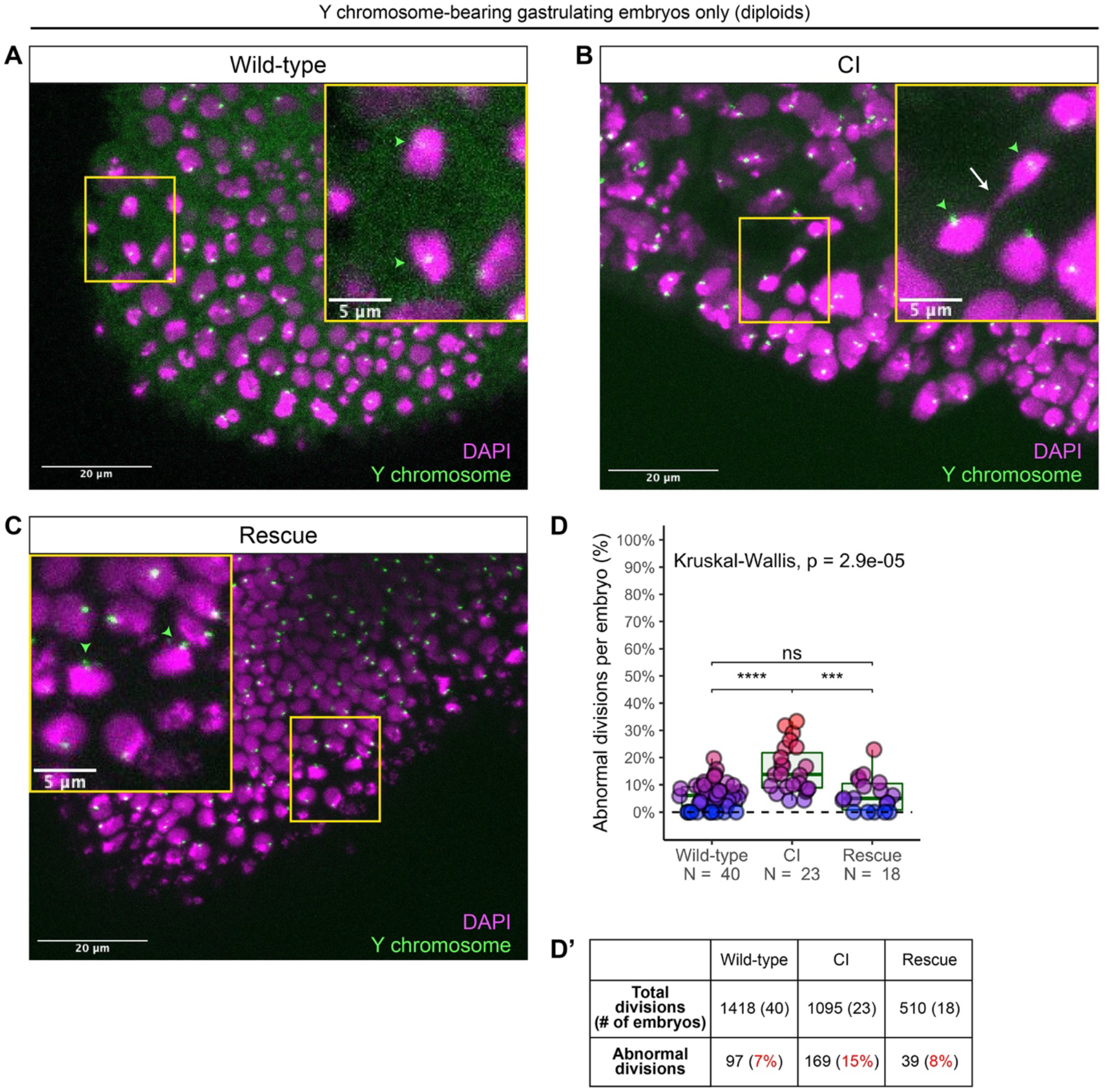
Diploid CI-derived gastrulating embryos that have escaped the first division defect exhibit increased chromosome segregation errors. (A-C) Gastrulating embryos from wild-type (A), CI (B), and rescue (C) crosses. Embryos are hybridized with probes that specifically recognize the *D. simulans* Y chromosome (green arrowheads) to select for diploidy (both maternal and paternal chromosome sets present) and counterstained with DAPI (magenta). Yellow boxes indicate zoomed in regions. Scale bars are 20 µm and 5 µm for unzoomed and zoomed images respectively. (A) Diploid wild-type-derived embryos exhibit relatively normal chromosome segregation. (B) Diploid CI-derived embryos have elevated rates of bridging and lagging chromosomes (arrow). (C) Diploid rescue-derived embryos exhibit relatively normal chromosome segregation. (D) Comparison of the percentage of chromosome segregation errors observed in diploid wild-type-, CI-, and rescue-derived embryos. Each dot represents one embryo. (D’) Summary of chromosome segregation errors in wild-type-, CI-, and rescue-derived embryos. *See also Figure S4*.

Although these CI-derived embryos are diploid and are likely to hatch despite the observed division defects, we found a subsequent decrease in the rate of hatched eggs that develop into adults in CI crosses compared to both wild-type and rescue crosses (Figure S4). Therefore, *Wolbachia* in the sperm may induce remarkably deferred CI defects that contribute to the selective advantage of infected females by promoting increased lethality during post-hatching development.

## DISCUSSION

In addition to the well-characterized early embryonic arrest, a number of reports reveal a large portion of CI-derived embryos undergo substantial embryonic development but then fail to hatch (Bonneau et al., 2018; Callaini et al., 1997; Callaini et al., 1996; Duron & Weill, 2006). This is in part explained by the behavior of the paternal chromosomes during the first division. While weak CI results in defective paternal chromosome segregation creating aneuploid nuclei that arrest in early embryonic development, strong CI results in complete failure of sister chromosome segregation and haploid nuclei bearing only the maternal chromosome complement (Bonneau et al., 2018; Callaini et al., 1997; Duron & Weill, 2006; Tram et al., 2006). Similar to diploid embryos that have “escaped” the first division defects, haploid embryos develop normally to cellular blastulation. However, haploids then fail prior to hatching. Here, we addressed whether defects observed in late CI embryos such as chromosome segregation errors and nuclear fallout are the result of first division errors or a second, distinct CI-induced defect. Specifically, we examined the timing, extent, and causes of defects produced in late *D. simulans* embryos derived from CI crosses.

In accord with previous studies in *D. simulans* (Callaini et al., 1997; Callaini et al., 1996), we found that there is a second late embryonic lethality associated with CI-derived embryos: between one-fourth to one-third of embryos die after cellularization but before hatching (Figure 1). In contrast to previous reports (Lassy & Karr, 1996), we found these embryos initially proceeded normally through nuclear cycles 2-9 (Figure 2). However, as the CI-derived embryos progressed through the cortical divisions (cycles 10-14), they begin to experience increasingly more severe defects. These late embryonic defects include lagging anaphase chromosomes and chromosome bridging, which directly result in nuclear fallout, and further chromosome bridging during gastrulation (Figures 3, S2). The normal progression of embryos through cycles 2-9 suggests paternal chromosome segregation was not partially defective in the first division. This is because paternal chromosome bridging in the first division would produce aneuploid daughter nuclei bearing chromosome fragments lacking telomeres. The lack of telomeres would result in detectable breakage-fusion-bridge cycles and amplifications of the aneuploidy in subsequent divisions (McClintock, 1941; Titen & Golic, 2008), which we did not observe. Our sequencing analysis of cellularized embryos (Figure 5) confirms that late-stage CI embryos did not experience partial chromosome loss during the first division. Thus, the mitotic defects first observed in cortical syncytial divisions are unlikely a consequence of CI-induced segmental aneuploidy following the first nuclear cycle.

Consequently, our single embryo PCR and whole genome sequencing of cellularized blastoderms containing a paternally-derived *egfp* transgene revealed late-stage CI embryos were either haploid or diploid (Figure 5). The percentage of diploid embryos closely corresponded to the percentage of embryos hatched, suggesting late embryonic lethality is associated with CI-induced haploidy in *D. simulans*, as previously reported (Callaini et al., 1997). Importantly, this experiment also demonstrated that the detection of paternally-derived chromosomes in that late-stage CI embryos could be used to distinguish between embryos that had experienced first division defects (haploid=only maternal chromosomes, no paternal chromosomes) and embryos that had not experienced any first division defects (diploid=both maternal and paternal chromosomes). As discussed below, this allowed us to uncouple CI-induced late embryo defects from first division defects.

In spite of the strong association between haploidy and lethality, first division-induced haploidy in and of itself cannot explain the defects we observed in CI-derived blastoderm and gastrulating embryos. This is because 1) haploidy is not intrinsically harmful to mitotic divisions in *Drosophila*. For example, in some *Drosophila* mutations that induce haploidy, chromosome segregation occurs normally during cortical divisions (Tang et al., 2017). Additionally, meiosis II—essentially a mitotic division of a haploid nucleus—is highly accurate by necessity. Furthermore, 2) any downstream effects of haploidy—such as changes to zygotic gene copy number or loss of zygotic heterozygosity—cannot explain defects first detected in syncytial cortical divisions (cycles 10-13), which do not require zygotic transcription (Yuan et al., 2016). In contrast, the observed defects in CI-derived late embryos are more likely due to a second, CI-induced defect.

In support of this hypothesis, our observation of increased chromosome segregation errors in diploid CI gastrulating embryos bearing paternally-derived Y chromosomes establishes that the defects observed in late-stage CI embryos are not limited to haploids (Figure 6). Instead, defects are present in diploid late-stage embryos. Significantly, as discussed above for the paternally-derived *egfp* transgene, detection of the Y chromosome by FISH allowed us to select late-stage diploid embryos that had “escaped” first division defects and instead continued development with both paternal and maternal chromosome complements. Therefore, the significant increase in mitotic errors observed in diploid CI-derived embryos relative to wild-type-derived embryos demonstrates the existence of a second, CI-induced defect, completely separate from the first division defect (Figure 7A-B). Significantly, maternally-supplied *Wolbachia* independently rescues this defect as well (Figure 7C).

**Figure 7.**
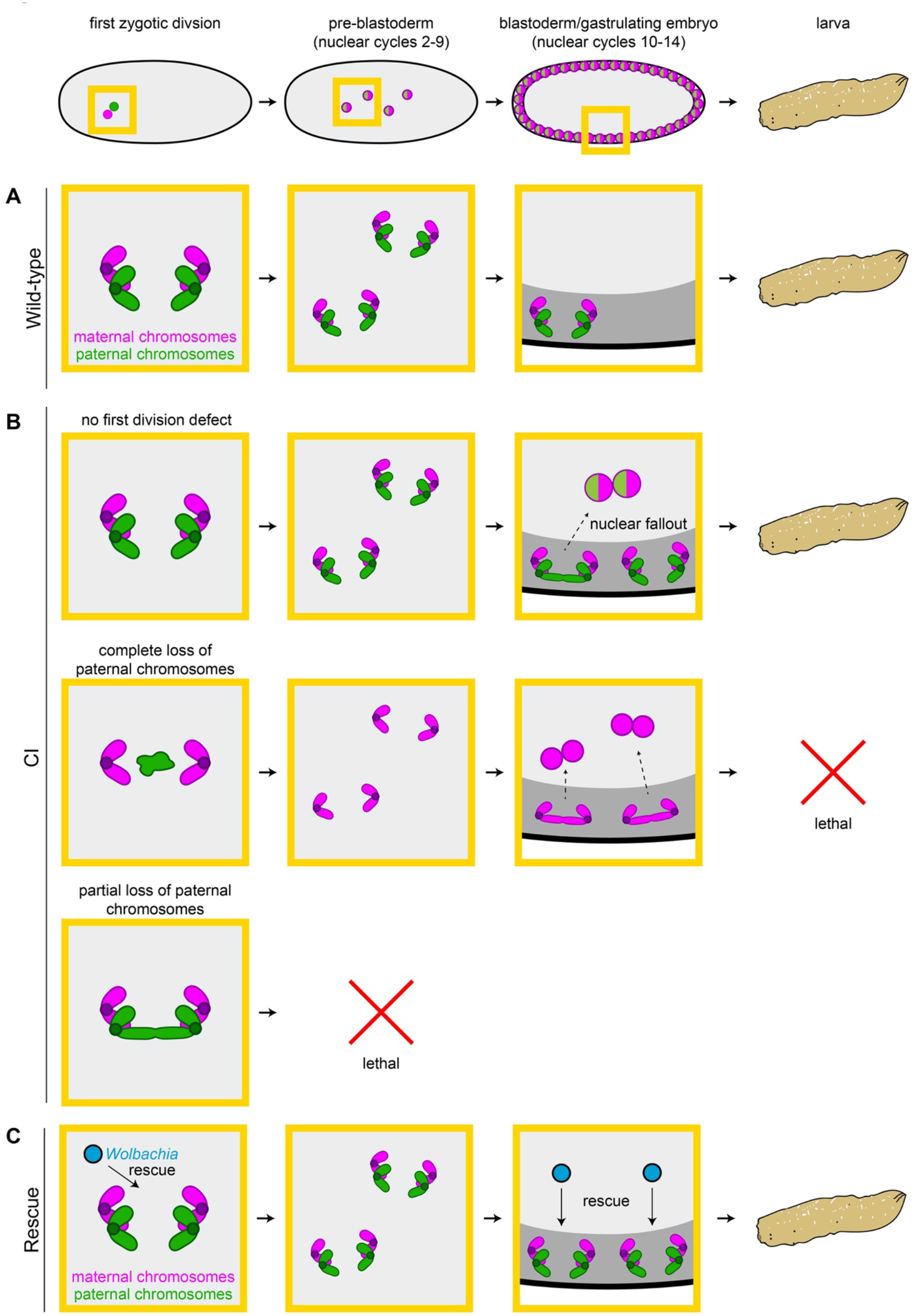
CI induces independent first division and mid-blastula transition chromosome segregation errors. (A) During the first zygotic division in wild-type derived embryos, paternal (green) and maternal (magenta) chromosomes segregate normally. Chromosome segregation occurs normally during pre-blastoderm, blastula, and post-cellularization divisions. Embryos hatch. (B, top row) In CI-derived embryos, if there are no segregation defects during the first division, embryos develop as diploids containing full maternal and paternal chromosome sets. Pre-blastoderm divisions proceed normally. However, during blastoderm stages, CI induces a second set of defects, which cause chromosome segregation errors and subsequent nuclear fallout (dashed arrow). Chromosome segregation errors continue during gastrulation. These defects occur at moderate frequencies and embryos hatch. (B, middle row) If the paternal chromosomes are completely excluded during the first division, embryos develop as haploids from only the maternal chromosome set. Pre-blastoderm divisions proceed normally, followed by increased chromosome segregation errors and nuclear fallout during blastoderm divisions. Chromosome segregation errors continue during gastrulation. Perhaps due to CI being strong in haploid embryos (Bonneau et al., 2018), this second set of CI-induced defects is more severe, and embryos fail to hatch, due to their haploidy. (B, bottom row) If the paternal chromosomes are partially lost during the first divisions, embryos arrest due to severe aneuploidy. (C) Maternally-supplied *Wolbachia* (blue circles) rescue both the first division defects and the late-stage defects independently.

Interestingly, we also observed several non-Y-bearing gastrulating embryos from CI crosses that had extensive chromosome segregation errors beyond what we had observed for the diploid Y-bearing gastrulating embryos (Figure S5A-C). Non-Y-bearing embryos may either be diploid (XX) or haploid (XØ). If these embryos were haploid, this observation would suggest that CI could affect both the maternal chromosomes and paternal chromosomes.

One intriguing aspect of the second CI-induced defect is that the embryos progress normally through the early mitotic cycles and then begin to exhibit mitotic defects in the blastoderm stage. The explanation is likely a consequence of the dramatic structural and regulator cell cycle modifications that occur when the dividing nuclei arrive at the cortex (Farrell & O’Farrell, 2014). These include heterochromatin formation, initiation of late origins of replication, slowing of DNA replication, activation of zygotic transcription, and metaphase furrow formation (Farrell & O’Farrell, 2014; Li et al., 2014; Riggs et al., 2003; Seller et al., 2019; Seller & O’Farrell, 2018). The phenotype of numerous maternal-effect mutations that either rely on or disrupt these processes is strikingly similar to the defects observed in CI embryos: normal early pre-cortical divisions followed by extensive mitotic errors and nuclear fallout during the cortical blastoderm divisions (Sullivan & Theurkauf, 1995). For example, because of the slowing of DNA replication during the cortical divisions, activation of the S-phase checkpoint is specifically required during this stage. Maternal-effect mutants that disrupt this checkpoint progress normally through the early divisions but exhibit anaphase bridging and nuclear fallout during the late cortical blastoderm divisions as a result of entering metaphase with incompletely replicated chromosomes (Fogarty et al., 1997; Fogarty et al., 1994). Given the similarity of this phenotype, both in timing and chromosome dynamics, CI-induced late division defects may be due to improper chromosome replication. Additionally, defects in other events specific to the cortical blastoderm cycles, may also contribute directly or indirectly to CI-induced defects. For example, studies of hybrid incompatibilities between *D. simulans* and *D. melanogaster* show that heterochromatin establishment may be particularly sensitive, and its disruption can result in defects strikingly similar to the late CI defects (Ferree & Barbash, 2009). Other important processes, such as those involved in DNA integrity, protein turnover, and cell cycle timing, may be also involved (Momtaz et al., 2020).

In considering the mechanism by which paternal *Wolbachia* may induce these defects, the observation that CI-derived blastoderm embryos progress normally through pre-cortical divisions must be noted. One potential explanation is that the chromosomes in CI-derived embryos could be epigenetically marked by *Wolbachia* in the paternal germline. This mark would then persist through the pre-cortical divisions and become disruptive during blastoderm divisions. Interestingly, *Wolbachia* infection results in altered DNA methylation levels in certain wasps, mosquitos, and *Drosophila* (LePage et al., 2014; Wu et al., 2020; Ye et al., 2013). Should *Wolbachia* in the male germline similarly change the low naturally occurring methylation levels in *D. simulans* (Deshmukh et al., 2018), the altered mark may become disruptive in blastoderm divisions, potentially by distorting heterochromatin establishment. However, DNA methylation does not appear to contribute to CI levels (LePage et al., 2014).

An alternative explanation for the specificity of the late blastoderm defects comes from studies into hybrid dysgenesis in *D. melanogaster* in which unregulated mobilization of transposable elements results in a spectrum of genetic and developmental defects in the germlines of dysgenic progeny (Kidwell et al., 1977). Transposition in progeny can be suppressed when maternally-supplied small RNAs mediate silencing of the transposable element (Czech & Hannon, 2016). This silencing is associated with increased H3K9 methylation, increased heterochromatin levels, and altered splicing (Le Thomas et al., 2013; Sienski et al., 2012; Teixeira et al., 2017). Given small RNAs can affect chromosomes in *trans* (Hermant et al., 2015), CI may induce a similar small RNA pathway that could epigenetically alter both paternal and maternal chromosomes prior to the first division. As the blastoderm divisions do not require zygotic transcription (Yuan et al., 2016), it is unlikely an epigenetic alteration, if occurring, would cause defects via disrupted transcription. Instead, as discussed above, an epigenetic change may disrupt key aspects of the mid-blastula transition, which in turn could result in the observed errors.

Insight into the molecular mechanism of CI came with the discovery of the *Wolbachia*-encoded Cifs that play a key role in CI and rescue (Beckmann et al., 2017; LePage et al., 2017). A combination of molecular, genetic, and biochemical studies provided compelling evidence that the *Wolbachia* encoded genes, *CidB* and *CidA*, act as a paternally supplied toxin and maternally supplied anti-toxin respectively (Beckmann et al., 2017; Horard et al., 2022; Wang et al., 2022). However, a toxin/anti-toxin model for CI does not easily explain the cortical blastoderm defects occurring after many rounds of normal mitotic cycles. This is because a paternally-supplied toxin is expected to be diluted with every round of division, and therefore its induced-defects would decrease over time. Similarly, a second set of CI/rescue elements, *CinA* and *CinB*, is also proposed to act in a toxin/anti-toxin manner (Chen et al., 2019; Sun et al., 2022). An alternative possibility is that Cifs may epigenetically modify paternal and maternal chromosomes to mediate CI and rescue (Kaur et al., 2022).

How and if these proteins also contribute to CI-induced late-embryo defects remains to be determined. Our observation of chromosome segregation errors in diploid embryos that have progressed normally through the first division (and thus should have minimal Cif activity) suggests an additional set of *Wolbachia* genes may induce late-embryo defects. Additionally, unlike the Cif-mediated first division errors, CI-induced mitotic defects in late embryos do not appear to result from abnormal condensation, alignment, or timing of metaphase exit (Figure 4), suggesting a separate proximate cause. Instead, the observed chromosome bridging is strikingly similar to embryos exposed to the DNA replication inhibitor aphidicolin (Farrell et al., 2012), suggesting CI-derived blastoderm embryos may be entering anaphase with incompletely replicated chromosomes. Thus, any model of CI and rescue, be it toxin/anti-toxin, lock-key, titration, or timing, must account for the fact that some effects of *Wolbachia* on the sperm are not realized until hours and many cell cycles later when the embryos progress through the mid-blastula transition and the late blastoderm divisions.

## MATERIALS AND METHODS

### Drosophila stocks

All stocks were grown on standard brown food (Sullivan, 2000) at 25°C with a 12h light/dark cycle. Uninfected *Drosophila simulans* stocks were generated by tetracycline-curing a *w*-*Wolbachia* (*wRiv*)-infected stock (Serbus et al., 2015). Uninfected and infected stocks were allowed to grow for many generations prior to their use. Throughout these experiments, we routinely checked for the presence/absence of *Wolbachia* by PCR with primers against the 16s rRNA gene of *Wolbachia*.

An uninfected *egfp*-bearing stock was obtained from the National *Drosophila* Species Stock Center (Cornell College of Agriculture and Life Sciences; #275; *w*[501]; PBac(GreenEye.UAS.tubEGFP)Dsim3) (Holtzman et al., 2010). *Wolbachia* was introduced to this stock by crossing to *Wolbachia*-infected females. Progeny was backcrossed to obtain flies homozygous for *egfp*. Stocks were routinely checked for *Wolbachia* and *egfp* presence by PCR with primers against the 16s rRNA gene of *Wolbachia* and *egfp* respectively. Males from this stock were used for experiments in which infected father transmitted an *egfp* transgene to offspring.

Embryos were collected from crosses of 3-5 day old flies (Figures 1, S1, 2, 3) or 2-4 day old flies (Figures S2, 4-6). For experiments in Figure 1-3 embryos were collected for 4 days after the initial collection. For all other experiments, embryos were collected only on the initial collection.

### Egg hatch assays

For experiments involving egg hatch assays (Figure 1, S1, 4), collected embryos were aged in a humid chamber at 25°C for at least 30h before hatched eggs were counted.

### Embryo fixation

For fixed experiments assaying embryo stage, abnormalities, and nuclear fallout, 1-6h (Figure 1A-B’), 2.5-3h (Figure 1C-D’), 0-4h (Figure 2), and 1-4h (Figures 3) embryos were dechorionated in bleach, washed thoroughly in water, and transferred to a 1:1 ratio of heptane and 32% paraformaldehyde for 5 min. Paraformaldehyde was subsequently removed and replaced with methanol and shaken vigorously. Heptane was removed and embryos stored in methanol at 4°C. Embryos were mounted directly in PI (Figure 1A-B’) or DAPI with Vectashield (Vector H-1200) (Figures 1C-D’, 2, 3).

For fixed experiments analyzing nuclear detachment from centrosomes (Figure 3), 1-4h embryos were initially fixed as described above. Embryos were rehydrated in PBT (PBS + 0.05% Triton + 1% BSA), blocked for 1h, and incubated with rabbit anti-centrosomin antibody (1:200) (Megraw et al., 1999). After 3x washes in PBT, embryos were incubated with anti-rabbit-Alexa488 secondary (1:1000 ThermoFisher A-11008). Embryos were washed 3x in PBT, rinsed 4X in PBS, and counterstained with DAPI in Vectashield.

For fixed experiments assaying chromosome segregation errors in gastrulating embryos (Figure S2), 3-5h embryos were dechorionated in bleach, washed thoroughly in water, and permeabilized in heptane for 2.5 min. Embryos were fixed by adding an equal volume methanol to the heptane and shaking vigorously. Heptane was removed, and embryos stored at 4°C in methanol. Embryos were mounted directly in DAPI with Vectashield.

For fixed experiments involving fluorescence in situ hybridization (Figure 6, S5), 2-5h embryos were dechorionated in bleach, washed thoroughly in water, and permeabilized in ice cold heptane for 2.5 min. Embryos were fixed in an ice cold 4% paraformaldehyde-46% PBS-50% heptane mixture for 10 min. Following removal of the paraformaldehyde-PBS solution, an equal volume of ice cold methanol was added to the heptane and shaken vigorously. Heptane was then removed. Embryos were then stored at 4°C in methanol.

### Live embryo staging

For experiments involving live embryo staging (Figures S1, 5), embryos were collected for 45 min, hand dechorionated, covered in halocarbon oil, and aged in a humid chamber at 25°C for 2.5h. Embryo stage was either scored after this time (Figure S1) or for every 60 min (Figure 5). Embryos were staged using an Olympus SZH10 high-powered dissecting scope. Live images presented in Figure 1 were acquired on a Zeiss Axiozoom V.16 microscope equipped with a Zeiss AxioCam HRm monochrome camera. Images were acquired with Zeiss Zen software.

### Embryo injection

For live imaging experiments (Figure 4), 0.5-1.5h embryos were hand dechorionated, placed in halocarbon oil, and injected with rhodamine-labeled histone. Embryos were imaged directly after injection in areas adjacent to the injection site.

### Fluorescence in situ hybridization

Alexa488-conjugated probes targeting the *D. simulans* Y-chromosome (AAT-AAA-C)_4_ (Ferree & Barbash, 2009) were synthesized by Integrated DNA Technologies (Coralville, IA, USA). Paraformaldehyde-fixed embryos were rehydrated in PBT (PBS + 0.05% Triton + 1% BSA). Embryos were washed in 4x saline sodium citrate (SSC), 10% formamide, 50mM imidazole for 1h at 37°C. Embryos were hybridized with probes in hybridization buffer (4x SSC, 10% formamide, 0.0001% dextran sulfate) at 92°C for 3 min then 37°C overnight. Embryos were washed 3x in 2x SSC, 50% formamide for 10 min at 37°C, rinsed 4x in PBS, and counterstained with DAPI in Vectashield.

### Confocal imaging

Live and fixed embryo imaging was performed on an inverted Leica DMI6000 SP5 scanning confocal microscope. DAPI was excited with a 405 nm laser and collected from 410-480 nm. Alexa488 was excited with a 488 nm laser and collected from 518-584 nm. Rhodamine was excited with a 543 nm laser and collected from 555-620 nm. PI was excited with 514 and 543 nm lasers and collected from 627-732 nm. Embryos were imaged with either 10x/0.3, 20x/0.75, 40x/1.25 oil, or 63x/1.4 oil objectives. All imaging was performed at room temperature. Images were acquired with Leica Application Suite Advanced Fluorescence software. For live imaging experiments (Figure 4), timepoints between images were every 12-60 sec depending on the size of the z-stack.

### Single embryo PCR analysis

Cellularized blastoderms were individually squashed and then lysed in 10 µL buffer containing Proteinase K and ThermoPol reaction buffer (New England BioLabs) for 45 min at 60°C then 10 min at 95°C. PCR was run with 1 µL sample in 20 µL total reaction volume, using primers targeting *egfp* (5’: ATCAAGCTTGTGAGCAAGGGCGAGGAGC; and 3’: ACCTCGAGCTACTTGTACAGCTCGTCCATGC) (Cruachem). PCR was run as: 10 min at 95°C, 31x(30 sec at 95°C, 1 min at 60°C, 1 min at 72°C), 10 min at 72°C. PCR products were resolved on a 1% agarose gel. These conditions regularly produced an ∼1.4kb band only when the *egfp* transgene was present.

### Single embryo sequencing

Cellularized blastoderms were individually squashed, frozen in liquid nitrogen, and stored at - 80°C. Library preparation (NexteraXT kit) and paired-end sequencing (Illumina HiSeq, 2×150bp) was performed by Azenta Life Sciences (Indianapolis, IN, USA). As samples contained host DNA, *Wolbachia* (*wRi*) DNA, and an *egfp* insertion, we assembled a reference genome consisting of the *D. simulans* genome (WUGSC mosaic 1.0/droSim1 assembly (Drosophila 12 Genomes et al., 2007), UCSC Genome Browser, Santa Cruz, CA, USA), a *wRI* genome ((Klasson et al., 2009) GenBank CP001391.1), and the *egfp* sequence from the *p-egfp* plasmid (Addgene, Watertown, MA, USA). We additionally included a 714 bp randomized sequence as a negative control.

Reads were aligned to the reference genome using BWA-MEM2 (2.2.1) (Md et al., 2019). Duplicate reads were removed using Picard tools (2.27.1) (http://broadinstitute.github.io/picard/) and low-quality reads (q<20) were subsequently removed. BEDTools (2.26.0) (Quinlan & Hall, 2010) was used to assign depth of coverage at each position in the genome. Read alignment and processing was performed using the Hummingbird Computational Cluster (UC Santa Cruz, Santa Cruz, CA, USA). Gene coordinate positions were determined in the UCSC Genome Browser.

Percent depth of a gene was calculated by dividing the average depth across a gene by the average depth across the whole genome for that embryo and multiplying by 100%. Embryos were considered diploid if the mean depth of reads aligning to the *egfp* transgene was meaningful (around 50% for heterozygote embryos) and reads were distributed evenly across the entirety of the *egfp* transgene. To decrease stochastic noise and accurately assess potential chromosome loss, we analyzed 5 genes from each chromosome/chromosome arm (Y, X, 2L, 2R, 3L, 3R, 4). Chromosome/chromosome arm loss was considered if the depth of reads across multiple genes on a chromosome/chromosome arm dropped from either 100% to 50% (diploid) or from 100% to 0% (haploid). As a proof of concept, an example of natural chromosome “loss” can be observed in male embryos (Y-linked genes present) in which the depth of reads mapping to genes on the X chromosome are ∼50% of the mean genome depth (hemizygous).

### Egg-to-adult assays

Egg hatch assays were performed using embryos collected from 2-4 day old flies. Eggs were counted and transferred to a new collection plate in a new collection bottle. Hatched eggs were counted after at least 30h. Adults were counted for each plate for as long as new adults were eclosing.

### Statistical analyses

All statistical analyses were performed in R (4.0.5, R core team). The following statistical tests were used: χ-square test (Figure 1, S1), two-sided paired t-test (Figure 1), two-sided Fisher’s exact test (Figure 2), Kruskal-Wallis test (Figures 3, S2, 6, S4), and Mann-Whitney tests (Figures S2, 4-6, S4).

### Figure preparation

Graphs were created in R using the ggplot2 package (Wickham, 2016). To improve the clarity of certain panels, images were adjusted for brightness and contrast in FIJI. Figures were assembled in Adobe Illustrator (Adobe, San Jose, CA, USA).

## ACKNOWLEDGEMENTS

We thank Dr. Benjamin Abrams (UCSC Life Sciences Microscopy Center, RRID: SCR_021135) for his technical support and assistance with microscopy experiments. We thank Dr. Shelbi Russell and Dr. Andreas Rechsteiner for their helpful advice on sequencing experiments and analysis. We thank Dr. Timothy Megraw who provided our lab with the anti-centrosomin antibody. We thank Dr. Jonathan Minden for providing rhodamine-labeled histones. This work was funded by an NIH grant NIGMS-1R35GM139595 awarded to W. Sullivan.

## AUTHOR CONTRIBUTIONS

W.S. conceived the manuscript. B.W. contributed to the planning, execution, and analysis of experiments involving live imaging (Figure 4), paired analysis of embryo stage and hatch rate (Figure 1), single embryo PCR and sequencing analysis (Figure 5, Figure S3), staining of gastrulating embryos (Figures S2, 6, S5), and egg-to-adult assays (Figure S4). S.T. contributed to the planning, execution, and analysis of experiments involving abnormal embryo counts (Figure 2), nuclear fallout (Figure 3), live imaging (Figure 4), and the planning of egg-to-adult assays (Figure S4). M.A. contributed the execution and analysis of experiments involving single embryo PCR and sequencing analysis (Figure 5, S3), staining of gastrulating embryos (Figures 6, S5), and egg-to-adult assays (Figure S4). G.V. contributed to the execution and analysis of experiments involving live imaging (Figure 4), paired analysis of embryo stage and hatch rate (Figure 1), and staining of gastrulating embryos (Figure S2). N.L. contributed to the execution and analysis of experiments involving embryo stage and hatch rate (Figures 1 and S1). K.H. contributed to the execution of paired analysis of embryo stage and hatch rate (Figure 1). W.S. contributed to the planning, execution, and analysis of experiments involving embryo stage and hatch rate (Figures 1 and S1), abnormal embryo counts (Figure 2), nuclear fallout (Figures 3), and live imaging (Figure 4). W.S., BW., S.T., and M.A. contributed to all aspects of manuscript writing and preparation.

## SUPPLEMENTAL INFORMATION

**Figure S1, related to Figure 1.**
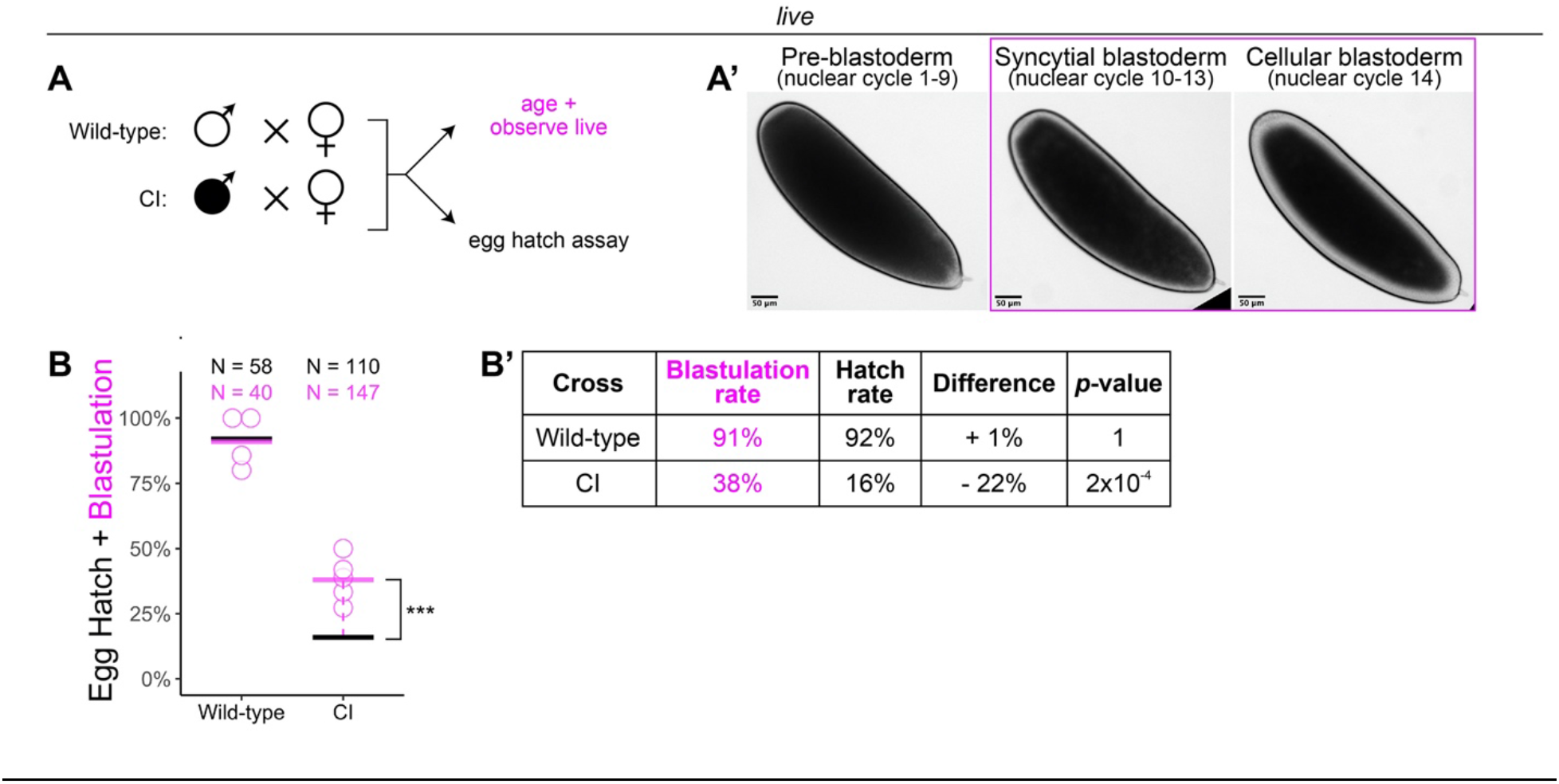
*Wolbachia* induces late embryonic lethality. (A) Embryos were collected from wild-type and CI crosses and either used for egg hatch assays or aged and observed live. (A’) Live observation of dechorionated embryos under a high-power dissecting scope enabled categorization of embryo stage as pre-blastoderm, syncytial blastoderm, or cellular blastoderms. Scale bars are 50 µm. (B-B’) Comparison between blastulation rate and egg hatch rate between wild-type and CI crosses. Each circle represents one round of live categorization. Black and magenta lines represent the egg hatch rate and the average blastulation rate respectively. See also Figure 1.

**Figure S2, related to Figure 3.**
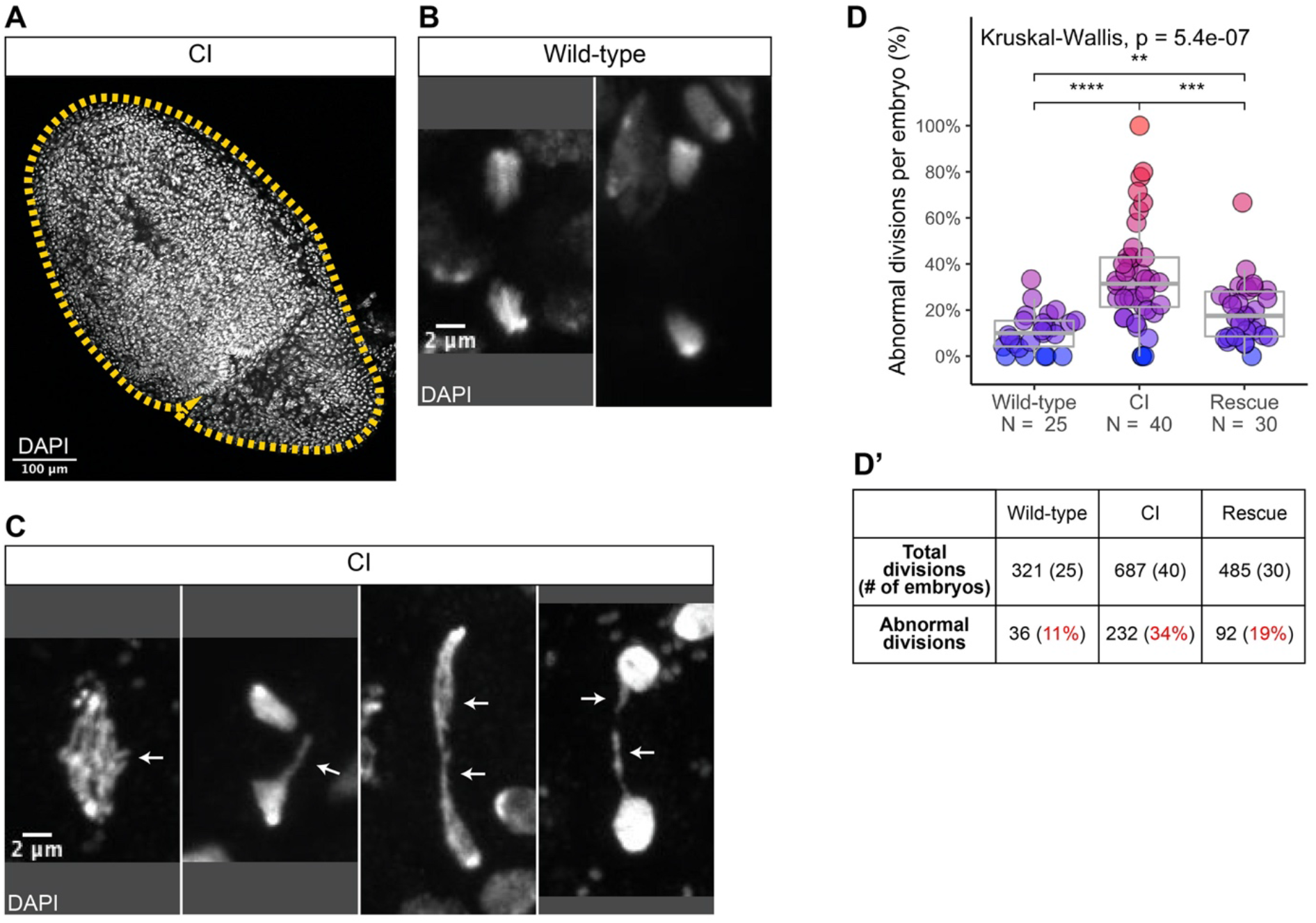
CI-derived embryos exhibit increased rates of chromosome segregation errors and nuclear fallout. (A) A whole CI-derived gastrulating embryo is outlined. Scale bars is 100 µm. (B-C) Examples of divisions observed in wild-type- (B) and CI- (C) derived gastrulating embryoss. While divisions from wild-type-derived embryos are normal, divisions from CI-derived embryos exhibit a variety of lagging/bridging chromosome segregation errors (arrows). Scale bars is 2 µm. (D-D’) Comparison of chromosome segregation errors in wild-type-, CI-, and rescue-derived gastrulating embryos. Each dot represents one embryo (D), and a summation is presented in (D’). See also Figure 3.

**Figure S3, related to Figure 5.**
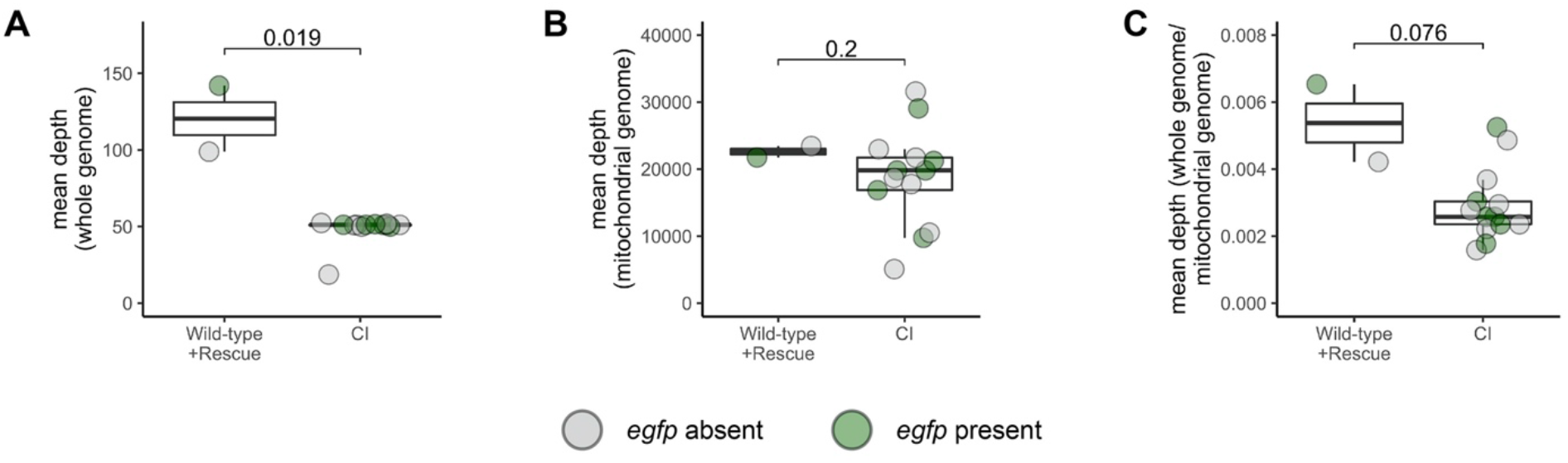
Depth of reads aligning to genome in CI-derived cellularized embryos is decreased compared to wild-type- and rescue-derived embryos. (A-C) Comparison of the mean depth of reads aligning to the reference genome (A), the mitochondrial genome (B), and the reference genome normalized to the mitochondrial genome (C) in wild-type-, CI-, and rescue-derived embryos. Each dot represents one embryo. Grey dots are embryos in which *egfp* was not detected (CI=haploids, wild-type=diploid). Green dots represent embryos in which *egfp* was detected (CI + Rescue = diploids). See also Figure 5.

**Figure S4, related to Figure 6.**
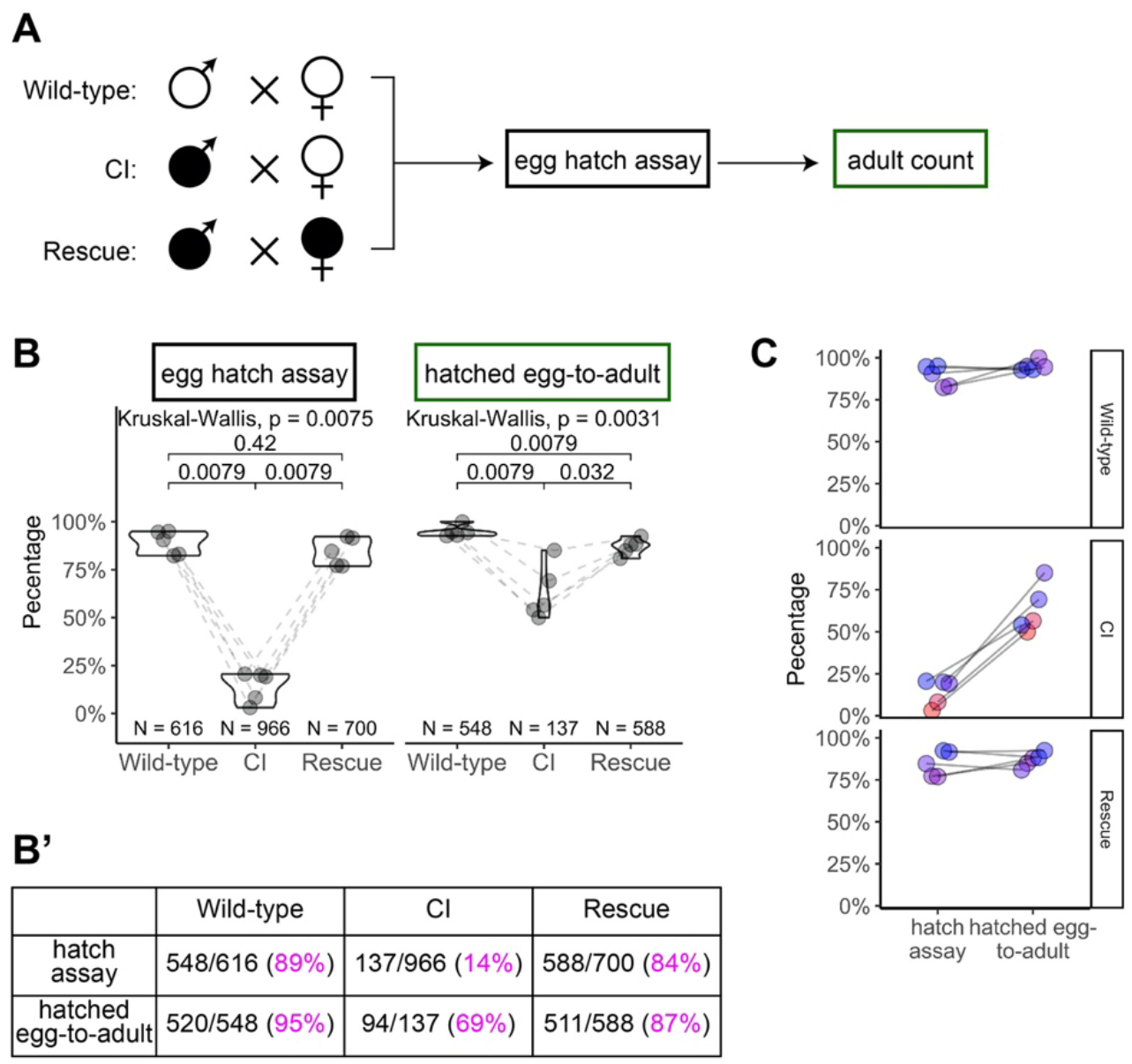
Hatched eggs from CI crosses exhibit significantly increased lethality prior to eclosion than those from wild-type or rescue crosses. (A) Diagram of the “egg-to-adult” experiment. Eggs from egg hatch assays were placed in new collection bottles. Adults were collected from hatched eggs and counted for as long as flies were eclosing. (B-B’) Comparison of egg hatch rates and hatched egg-to-adult rates. Hatched eggs from CI crosses had significantly reduced rates of developing into adults, suggesting the existence of a CI-induced lethal phase during larval development. Maternally-supplied Wolbachia rescues this larval lethality. Each dot represents one experiment/collection. Lines connect experiments performed simultaneously. Unless otherwise indicated, *p* values displayed were determined with Mann-Whitney tests. (C) Comparison of the strengths of early CI defects (as judged by hatching) and late CI defects (as judged by the hatched egg-to-adult rates). In general, higher lethality in hatching correlated to higher lethality during egg-to-adult development. Each dot represents one experiment. Lines connect the egg hatch rate and the hatched egg-to-adult rate for each experiment. See also Figure 6.

**Figure S5, related to Figure 6.**
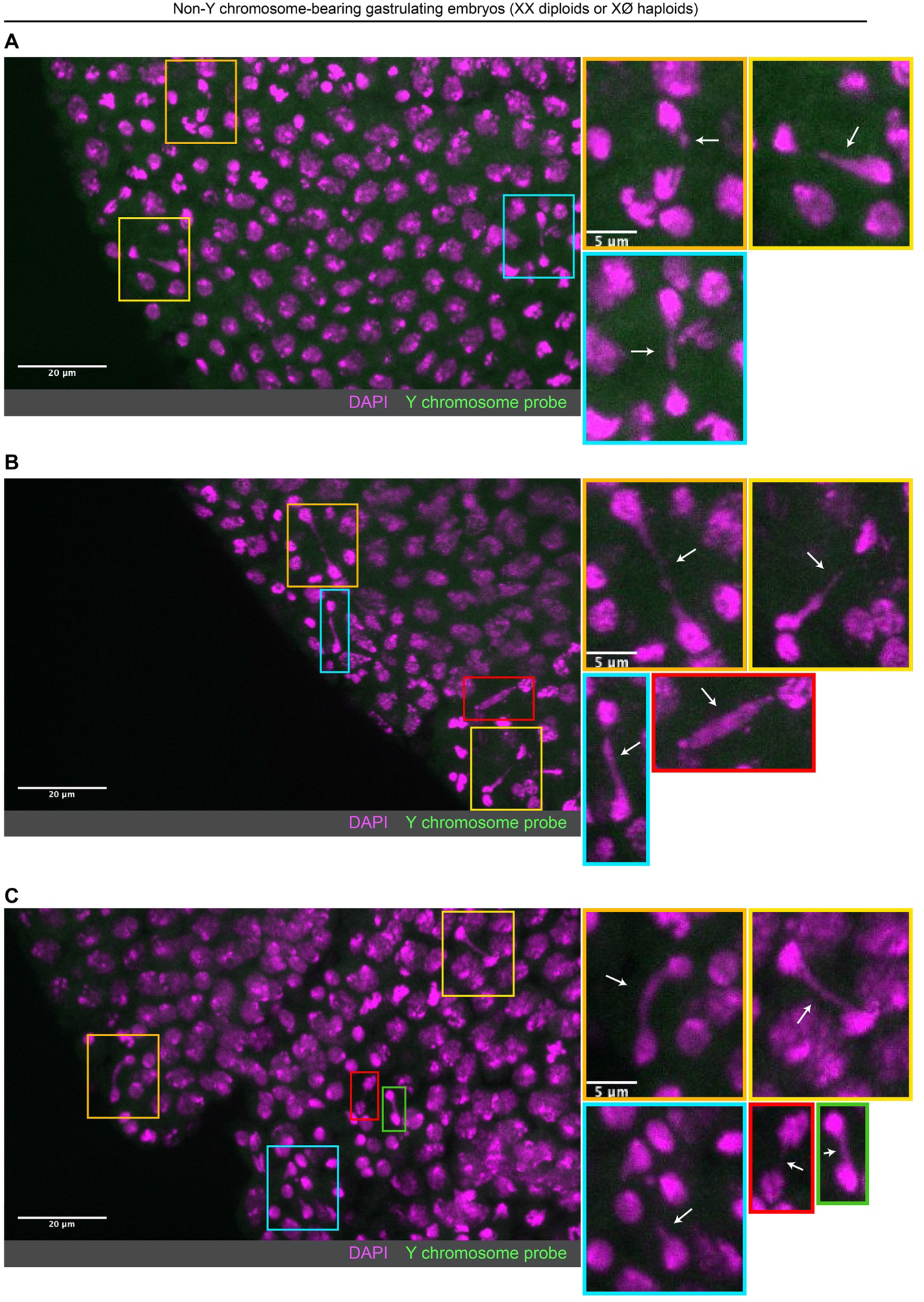
Non-Y chromosome-containing gastrulating embryos exhibit extensive chromosome segregation errors. (A-C) Examples of non-Y chromosome-containing gastrulating embryos (haploid or diploid) with extensive chromosome segregation errors. No probes targeting the Y chromosome (green) were detected in these embryos. Embryos were counterstained with DAPI (magenta). The extent of errors observed in these embryos is not observed in Y chromosome-containing diploid embryos, suggesting these embryos may be haploid and more strongly experience the second set of CI-induced defects. Boxes indicate zoomed in regions. Scale bars are 20 µm and 5 µm for unzoomed and zoomed images respectively. See also Figure 6.

## MOVIE LEGENDS

<u>Movie 1, related to Figure 4. Chromosome segregation immediately precedes nuclear fallout of cortical blastoderm nuclei</u>

A CI-derived embryo injected with rhodamine-labeled histone. Scale bar is 5 µm and time is in sec. See also Figure 4.

## SUPPLEMENTAL FILES

<u>File S1. Depth of coverage for coding sequences and *egfp* in wild-type-, CI-, and rescue-derived cellular blastoderms</u>

## REFERENCES

Beckmann, J. F., Ronau, J. A., & Hochstrasser, M. (2017). A Wolbachia deubiquitylating enzyme induces cytoplasmic incompatibility. Nat Microbiol, 2, 17007. https://doi.org/10.1038/nmicrobiol.2017.7

Bonneau, M., Atyame, C., Beji, M., Justy, F., Cohen-Gonsaud, M., Sicard, M., & Weill, M. (2018). Culex pipiens crossing type diversity is governed by an amplified and polymorphic operon of Wolbachia. Nat Commun, 9(1), 319. https://doi.org/10.1038/s41467-017-02749-w

Breeuwer, J. A., & Werren, J. H. (1990). Microorganisms associated with chromosome destruction and reproductive isolation between two insect species. Nature, 346(6284), 558–560. https://doi.org/10.1038/346558a0

Callaini, G., Dallai, R., & Riparbelli, M. G. (1997). Wolbachia-induced delay of paternal chromatin condensation does not prevent maternal chromosomes from entering anaphase in incompatible crosses of Drosophila simulans. J Cell Sci, 110 ( *Pt 2*), 271–280. https://doi.org/10.1242/jcs.110.2.271

Callaini, G., Riparbelli, M. G., Giordano, R., & Dallai, R. (1996). Mitotic defects associated with cytoplasmic incompatibility in Drosophila simulans. Journal of Invertebrate Pathology, 67(1), 55–64. https://doi.org/DOI 10.1006/jipa.1996.0009

Chen, H., Ronau, J. A., Beckmann, J. F., & Hochstrasser, M. (2019). A Wolbachia nuclease and its binding partner provide a distinct mechanism for cytoplasmic incompatibility. Proc Natl Acad Sci U S A, 116(44), 22314–22321. https://doi.org/10.1073/pnas.1914571116

Chen, H., Zhang, M., & Hochstrasser, M. (2020). The Biochemistry of Cytoplasmic Incompatibility Caused by Endosymbiotic Bacteria. Genes (Basel), 11(8). https://doi.org/10.3390/genes11080852

Czech, B., & Hannon, G. J. (2016). One Loop to Rule Them All: The Ping-Pong Cycle and piRNA-Guided Silencing. Trends Biochem Sci, 41(4), 324–337. https://doi.org/10.1016/j.tibs.2015.12.008

Debec, A. (1978). Haploid cell cultures of Drosophila melanogaster. Nature, 274(5668), 255–256. https://doi.org/10.1038/274255a0

Deshmukh, S., Ponnaluri, V. C., Dai, N., Pradhan, S., & Deobagkar, D. (2018). Levels of DNA cytosine methylation in the Drosophila genome. PeerJ, 6, e5119. https://doi.org/10.7717/peerj.5119

Drosophila 12 Genomes, C., Clark, A. G., Eisen, M. B., Smith, D. R., Bergman, C. M., Oliver, B., Markow, T. A., Kaufman, T. C., Kellis, M., Gelbart, W., Iyer, V. N., Pollard, D. A., Sackton, T. B., Larracuente, A. M., Singh, N. D., Abad, J. P., Abt, D. N., Adryan, B., Aguade, M., Akashi, H., Anderson, W. W., Aquadro, C. F., Ardell, D. H., Arguello, R., Artieri, C. G., Barbash, D. A., Barker, D., Barsanti, P., Batterham, P., Batzoglou, S., Begun, D., Bhutkar, A., Blanco, E., Bosak, S. A., Bradley, R. K., Brand, A. D., Brent, M. R., Brooks, A. N., Brown, R. H., Butlin, R. K., Caggese, C., Calvi, B. R., Bernardo de Carvalho, A., Caspi, A., Castrezana, S., Celniker, S. E., Chang, J. L., Chapple, C., Chatterji, S., Chinwalla, A., Civetta, A., Clifton, S. W., Comeron, J. M., Costello, J. C., Coyne, J. A., Daub, J., David, R. G., Delcher, A. L., Delehaunty, K., Do, C. B., Ebling, H., Edwards, K., Eickbush, T., Evans, J. D., Filipski, A., Findeiss, S., Freyhult, E., Fulton, L., Fulton, R., Garcia, A. C., Gardiner, A., Garfield, D. A., Garvin, B. E., Gibson, G., Gilbert, D., Gnerre, S., Godfrey, J., Good, R., Gotea, V., Gravely, B., Greenberg, A. J., Griffiths-Jones, S., Gross, S., Guigo, R., Gustafson, E. A., Haerty, W., Hahn, M. W., Halligan, D. L., Halpern, A. L., Halter, G. M., Han, M. V., Heger, A., Hillier, L., Hinrichs, A. S., Holmes, I., Hoskins, R. A., Hubisz, M. J., Hultmark, D., Huntley, M. A., Jaffe, D. B., Jagadeeshan, S., Jeck, W. R., Johnson, J., Jones, C. D., Jordan, W. C., Karpen, G. H., Kataoka, E., Keightley, P. D., Kheradpour, P., Kirkness, E. F., Koerich, L. B., Kristiansen, K., Kudrna, D., Kulathinal, R. J., Kumar, S., Kwok, R., Lander, E., Langley, C. H., Lapoint, R., Lazzaro, B. P., Lee, S. J., Levesque, L., Li, R., Lin, C. F., Lin, M. F., Lindblad-Toh, K., Llopart, A., Long, M., Low, L., Lozovsky, E., Lu, J., Luo, M., Machado, C. A., Makalowski, W., Marzo, M., Matsuda, M., Matzkin, L., McAllister, B., McBride, C. S., McKernan, B., McKernan, K., Mendez-Lago, M., Minx, P., Mollenhauer, M. U., Montooth, K., Mount, S. M., Mu, X., Myers, E., Negre, B., Newfeld, S., Nielsen, R., Noor, M. A., O’Grady, P., Pachter, L., Papaceit, M., Parisi, M. J., Parisi, M., Parts, L., Pedersen, J. S., Pesole, G., Phillippy, A. M., Ponting, C. P., Pop, M., Porcelli, D., Powell, J. R., Prohaska, S., Pruitt, K., Puig, M., Quesneville, H., Ram, K. R., Rand, D., Rasmussen, M. D., Reed, L. K., Reenan, R., Reily, A., Remington, K. A., Rieger, T. T., Ritchie, M. G., Robin, C., Rogers, Y. H., Rohde, C., Rozas, J., Rubenfield, M. J., Ruiz, A., Russo, S., Salzberg, S. L., Sanchez-Gracia, A., Saranga, D. J., Sato, H., Schaeffer, S. W., Schatz, M. C., Schlenke, T., Schwartz, R., Segarra, C., Singh, R. S., Sirot, L., Sirota, M., Sisneros, N. B., Smith, C. D., Smith, T. F., Spieth, J., Stage, D. E., Stark, A., Stephan, W., Strausberg, R. L., Strempel, S., Sturgill, D., Sutton, G., Sutton, G. G., Tao, W., Teichmann, S., Tobari, Y. N., Tomimura, Y., Tsolas, J. M., Valente, V. L., Venter, E., Venter, J. C., Vicario, S., Vieira, F. G., Vilella, A. J., Villasante, A., Walenz, B., Wang, J., Wasserman, M., Watts, T., Wilson, D., Wilson, R. K., Wing, R. A., Wolfner, M. F., Wong, A., Wong, G. K., Wu, C. I., Wu, G., Yamamoto, D., Yang, H. P., Yang, S. P., Yorke, J. A., Yoshida, K., Zdobnov, E., Zhang, P., Zhang, Y., Zimin, A. V., Baldwin, J., Abdouelleil, A., Abdulkadir, J., Abebe, A., Abera, B., Abreu, J., Acer, S. C., Aftuck, L., Alexander, A., An, P., Anderson, E., Anderson, S., Arachi, H., Azer, M., Bachantsang, P., Barry, A., Bayul, T., Berlin, A., Bessette, D., Bloom, T., Blye, J., Boguslavskiy, L., Bonnet, C., Boukhgalter, B., Bourzgui, I., Brown, A., Cahill, P., Channer, S., Cheshatsang, Y., Chuda, L., Citroen, M., Collymore, A., Cooke, P., Costello, M., D’Aco, K., Daza, R., De Haan, G., DeGray, S., DeMaso, C., Dhargay, N., Dooley, K., Dooley, E., Doricent, M., Dorje, P., Dorjee, K., Dupes, A., Elong, R., Falk, J., Farina, A., Faro, S., Ferguson, D., Fisher, S., Foley, C. D., Franke, A., Friedrich, D., Gadbois, L., Gearin, G., Gearin, C. R., Giannoukos, G., Goode, T., Graham, J., Grandbois, E., Grewal, S., Gyaltsen, K., Hafez, N., Hagos, B., Hall, J., Henson, C., Hollinger, A., Honan, T., Huard, M. D., Hughes, L., Hurhula, B., Husby, M. E., Kamat, A., Kanga, B., Kashin, S., Khazanovich, D., Kisner, P., Lance, K., Lara, M., Lee, W., Lennon, N., Letendre, F., LeVine, R., Lipovsky, A., Liu, X., Liu, J., Liu, S., Lokyitsang, T., Lokyitsang, Y., Lubonja, R., Lui, A., MacDonald, P., Magnisalis, V., Maru, K., Matthews, C., McCusker, W., McDonough, S., Mehta, T., Meldrim, J., Meneus, L., Mihai, O., Mihalev, A., Mihova, T., Mittelman, R., Mlenga, V., Montmayeur, A., Mulrain, L., Navidi, A., Naylor, J., Negash, T., Nguyen, T., Nguyen, N., Nicol, R., Norbu, C., Norbu, N., Novod, N., O’Neill, B., Osman, S., Markiewicz, E., Oyono, O. L., Patti, C., Phunkhang, P., Pierre, F., Priest, M., Raghuraman, S., Rege, F., Reyes, R., Rise, C., Rogov, P., Ross, K., Ryan, E., Settipalli, S., Shea, T., Sherpa, N., Shi, L., Shih, D., Sparrow, T., Spaulding, J., Stalker, J., Stange-Thomann, N., Stavropoulos, S., Stone, C., Strader, C., Tesfaye, S., Thomson, T., Thoulutsang, Y., Thoulutsang, D., Topham, K., Topping, I., Tsamla, T., Vassiliev, H., Vo, A., Wangchuk, T., Wangdi, T., Weiand, M., Wilkinson, J., Wilson, A., Yadav, S., Young, G., Yu, Q., Zembek, L., Zhong, D., Zimmer, A., Zwirko, Z., Jaffe, D. B., Alvarez, P., Brockman, W., Butler, J., Chin, C., Gnerre, S., Grabherr, M., Kleber, M., Mauceli, E., & MacCallum, I. (2007). Evolution of genes and genomes on the Drosophila phylogeny. Nature, 450(7167), 203–218. https://doi.org/10.1038/nature06341

Duron, O., & Weill, M. (2006). Wolbachia infection influences the development of Culex pipiens embryo in incompatible crosses. Heredity (Edinb), 96(6), 493–500. https://doi.org/10.1038/sj.hdy.6800831

Farrell, J. A., & O’Farrell, P. H. (2014). From egg to gastrula: how the cell cycle is remodeled during the Drosophila mid-blastula transition. Annu Rev Genet, 48, 269–294. https://doi.org/10.1146/annurev-genet-111212-133531

Farrell, J. A., Shermoen, A. W., Yuan, K., & O’Farrell, P. H. (2012). Embryonic onset of late replication requires Cdc25 down-regulation. Genes Dev, 26(7), 714–725. https://doi.org/10.1101/gad.186429.111

Ferree, P. M., & Barbash, D. A. (2009). Species-specific heterochromatin prevents mitotic chromosome segregation to cause hybrid lethality in Drosophila. PLoS Biol, 7(10), e1000234. https://doi.org/10.1371/journal.pbio.1000234

Foe, V. E. (1989). Mitotic domains reveal early commitment of cells in Drosophila embryos. Development, 107(1), 1–22. https://www.ncbi.nlm.nih.gov/pubmed/2516798

Fogarty, P., Campbell, S. D., Abu-Shumays, R., Phalle, B. S., Yu, K. R., Uy, G. L., Goldberg, M. L., & Sullivan, W. (1997). The Drosophila grapes gene is related to checkpoint gene chk1/rad27 and is required for late syncytial division fidelity. Curr Biol, 7(6), 418–426. https://doi.org/10.1016/s0960-9822(06)00189-8

Fogarty, P., Kalpin, R. F., & Sullivan, W. (1994). The Drosophila maternal-effect mutation grapes causes a metaphase arrest at nuclear cycle 13. Development, 120(8), 2131–2142. https://doi.org/10.1242/dev.120.8.2131

Ghelelovitch, S. (1952). [Genetic determinism of sterility in the cross-breeding of various strains of Culex autogenicus Roubaud]. C R Hebd Seances Acad Sci, 234(24), 2386–2388. https://www.ncbi.nlm.nih.gov/pubmed/12979357 (Sur le determinisme genetique de la sterilite dans les croisements entre differentes souches de Culex autogenicus Roubaud.)

Hermant, C., Boivin, A., Teysset, L., Delmarre, V., Asif-Laidin, A., van den Beek, M., Antoniewski, C., & Ronsseray, S. (2015). Paramutation in Drosophila Requires Both Nuclear and Cytoplasmic Actors of the piRNA Pathway and Induces Cis-spreading of piRNA Production. Genetics, 201(4), 1381–1396. https://doi.org/10.1534/genetics.115.180307

Hoffmann, A. A., Turelli, M., & Simmons, G. M. (1986). Unidirectional Incompatibility between Populations of Drosophila Simulans. Evolution, 40(4), 692–701. https://doi.org/10.1111/j.1558-5646.1986.tb00531.x

Holtzman, S., Miller, D., Eisman, R., Kuwayama, H., Niimi, T., & Kaufman, T. (2010). Transgenic tools for members of the genus Drosophila with sequenced genomes. Fly (Austin), 4(4), 349–362. https://doi.org/10.4161/fly.4.4.13304

Horard, B., Terretaz, K., Gosselin-Grenet, A. S., Sobry, H., Sicard, M., Landmann, F., & Loppin, B. (2022). Paternal transmission of the Wolbachia CidB toxin underlies cytoplasmic incompatibility. Curr Biol, 32(6), 1319–1331 e1315. https://doi.org/10.1016/j.cub.2022.01.052

Illmensee, K., & Mahowald, A. P. (1974). Transplantation of posterior polar plasm in Drosophila. Induction of germ cells at the anterior pole of the egg. Proc Natl Acad Sci U S A, 71(4), 1016–1020. https://doi.org/10.1073/pnas.71.4.1016

Jiggins, F. M. (2017). The spread of Wolbachia through mosquito populations. PLoS Biol, 15(6), e2002780. https://doi.org/10.1371/journal.pbio.2002780

Jost, E. (1970). [Genetic investigations on the incompatibility in the Culex pipiens complex]. Theor Appl Genet, 40(6), 251–256. https://doi.org/10.1007/BF00282034 (Genetische Untersuchungen zur Inkompatibilitat im Culex-pipiens-Komplex.)

Kaur, R., Leigh, B. A., Ritchie, I. T., & Bordenstein, S. R. (2022). The Cif proteins from Wolbachia prophage WO modify sperm genome integrity to establish cytoplasmic incompatibility. PLoS Biol, 20(5), e3001584. https://doi.org/10.1371/journal.pbio.3001584

Kaur, R., Shropshire, J. D., Cross, K. L., Leigh, B., Mansueto, A. J., Stewart, V., Bordenstein, S. R., & Bordenstein, S. R. (2021). Living in the endosymbiotic world of Wolbachia: A centennial review. Cell Host Microbe, 29(6), 879–893. https://doi.org/10.1016/j.chom.2021.03.006

Kidwell, M. G., Kidwell, J. F., & Sved, J. A. (1977). Hybrid Dysgenesis in DROSOPHILA MELANOGASTER: A Syndrome of Aberrant Traits Including Mutation, Sterility and Male Recombination. Genetics, 86(4), 813–833. https://doi.org/10.1093/genetics/86.4.813

Klasson, L., Westberg, J., Sapountzis, P., Naslund, K., Lutnaes, Y., Darby, A. C., Veneti, Z., Chen, L., Braig, H. R., Garrett, R., Bourtzis, K., & Andersson, S. G. (2009). The mosaic genome structure of the Wolbachia wRi strain infecting Drosophila simulans. Proc Natl Acad Sci U S A, 106(14), 5725–5730. https://doi.org/10.1073/pnas.0810753106

Landmann, F., Orsi, G. A., Loppin, B., & Sullivan, W. (2009). Wolbachia-mediated cytoplasmic incompatibility is associated with impaired histone deposition in the male pronucleus. PLoS Pathog, 5(3), e1000343. https://doi.org/10.1371/journal.ppat.1000343

Lassy, C. W., & Karr, T. L. (1996). Cytological analysis of fertilization and early embryonic development in incompatible crosses of Drosophila simulans. Mech Dev, 57(1), 47–58. https://doi.org/10.1016/0925-4773(96)00527-8

Le Thomas, A., Rogers, A. K., Webster, A., Marinov, G. K., Liao, S. E., Perkins, E. M., Hur, J. K., Aravin, A. A., & Toth, K. F. (2013). Piwi induces piRNA-guided transcriptional silencing and establishment of a repressive chromatin state. Genes Dev, 27(4), 390–399. https://doi.org/10.1101/gad.209841.112

LePage, D. P., Jernigan, K. K., & Bordenstein, S. R. (2014). The relative importance of DNA methylation and Dnmt2-mediated epigenetic regulation on Wolbachia densities and cytoplasmic incompatibility. PeerJ, 2, e678. https://doi.org/10.7717/peerj.678

LePage, D. P., Metcalf, J. A., Bordenstein, S. R., On, J., Perlmutter, J. I., Shropshire, J. D., Layton, E. M., Funkhouser-Jones, L. J., Beckmann, J. F., & Bordenstein, S. R. (2017). Prophage WO genes recapitulate and enhance Wolbachia-induced cytoplasmic incompatibility. Nature, 543(7644), 243–247. https://doi.org/10.1038/nature21391

Li, X. Y., Harrison, M. M., Villalta, J. E., Kaplan, T., & Eisen, M. B. (2014). Establishment of regions of genomic activity during the Drosophila maternal to zygotic transition. Elife, 3. https://doi.org/10.7554/eLife.03737

McClintock, B. (1941). The Stability of Broken Ends of Chromosomes in Zea Mays. Genetics, 26(2), 234–282. https://doi.org/10.1093/genetics/26.2.234

Md, V., Misra, S., Li, H., & Aluru, S. (2019). Efficient Architecture-Aware Acceleration of BWA-MEM for Multicore Systems. arXiv:1907.12931. Retrieved July 01, 2019, from https://ui.adsabs.harvard.edu/abs/2019arXiv190712931M

Megraw, T. L., Li, K., Kao, L. R., & Kaufman, T. C. (1999). The centrosomin protein is required for centrosome assembly and function during cleavage in Drosophila. Development, 126(13), 2829–2839. https://doi.org/10.1242/dev.126.13.2829

Momtaz, A. Z., Ahumada Sabagh, A. D., Gonzalez Amortegui, J. G., Salazar, S. A., Finessi, A., Hernandez, J., Christensen, S., & Serbus, L. R. (2020). A Role for Maternal Factors in Suppressing Cytoplasmic Incompatibility. Front Microbiol, 11, 576844. https://doi.org/10.3389/fmicb.2020.576844

Moretti, R., Yen, P. S., Houe, V., Lampazzi, E., Desiderio, A., Failloux, A. B., & Calvitti, M. (2018). Combining Wolbachia-induced sterility and virus protection to fight Aedes albopictus-borne viruses. PLoS Negl Trop Dis, 12(7), e0006626. https://doi.org/10.1371/journal.pntd.0006626

Quinlan, A. R., & Hall, I. M. (2010). BEDTools: a flexible suite of utilities for comparing genomic features. Bioinformatics, 26(6), 841–842. https://doi.org/10.1093/bioinformatics/btq033

Reed, K. M., & Werren, J. H. (1995). Induction of paternal genome loss by the paternal-sex-ratio chromosome and cytoplasmic incompatibility bacteria (Wolbachia): a comparative study of early embryonic events. Mol Reprod Dev, 40(4), 408–418. https://doi.org/10.1002/mrd.1080400404

Riggs, B., Rothwell, W., Mische, S., Hickson, G. R., Matheson, J., Hays, T. S., Gould, G. W., & Sullivan, W. (2003). Actin cytoskeleton remodeling during early Drosophila furrow formation requires recycling endosomal components Nuclear-fallout and Rab11. J Cell Biol, 163(1), 143–154. https://doi.org/10.1083/jcb.200305115

Ryan, S. L., & Saul, G. B., 2nd. (1968). Post-fertilization effect of incompatibility factors in Mormoniella. Mol Gen Genet, 103(1), 29–36. https://doi.org/10.1007/BF00271154

Seller, C. A., Cho, C. Y., & O’Farrell, P. H. (2019). Rapid embryonic cell cycles defer the establishment of heterochromatin by Eggless/SetDB1 in Drosophila. Genes Dev, 33(7-8), 403–417. https://doi.org/10.1101/gad.321646.118

Seller, C. A., & O’Farrell, P. H. (2018). Rif1 prolongs the embryonic S phase at the Drosophila mid-blastula transition. PLoS Biol, 16(5), e2005687. https://doi.org/10.1371/journal.pbio.2005687

Serbus, L. R., Casper-Lindley, C., Landmann, F., & Sullivan, W. (2008). The genetics and cell biology of Wolbachia-host interactions. Annu Rev Genet, 42, 683–707. https://doi.org/10.1146/annurev.genet.41.110306.130354

Serbus, L. R., White, P. M., Silva, J. P., Rabe, A., Teixeira, L., Albertson, R., & Sullivan, W. (2015). The impact of host diet on Wolbachia titer in Drosophila. PLoS Pathog, 11(3), e1004777. https://doi.org/10.1371/journal.ppat.1004777

Shropshire, J. D., Leigh, B., & Bordenstein, S. R. (2020). Symbiont-mediated cytoplasmic incompatibility: what have we learned in 50 years? Elife, 9. https://doi.org/10.7554/eLife.61989

Sienski, G., Donertas, D., & Brennecke, J. (2012). Transcriptional silencing of transposons by Piwi and maelstrom and its impact on chromatin state and gene expression. Cell, 151(5), 964–980. https://doi.org/10.1016/j.cell.2012.10.040

Sokac, A. M., Biel, N., & De Renzis, S. (2022). Membrane-actin interactions in morphogenesis: Lessons learned from Drosophila cellularization. Semin Cell Dev Biol. https://doi.org/10.1016/j.semcdb.2022.03.028

Sullivan, W., Ashburner, A., Hawley, R.S. (2000). Drosophila Protocols. Cold Spring Harbor Laboratory Press.

Sullivan, W., Daily, D. R., Fogarty, P., Yook, K. J., & Pimpinelli, S. (1993). Delays in anaphase initiation occur in individual nuclei of the syncytial Drosophila embryo. Mol Biol Cell, 4(9), 885–896. https://doi.org/10.1091/mbc.4.9.885

Sullivan, W., Minden, J. S., & Alberts, B. M. (1990). daughterless-abo-like, a Drosophila maternal-effect mutation that exhibits abnormal centrosome separation during the late blastoderm divisions. Development, 110(2), 311–323. https://doi.org/10.1242/dev.110.2.311

Sullivan, W., & Theurkauf, W. E. (1995). The cytoskeleton and morphogenesis of the early Drosophila embryo. Curr Opin Cell Biol, 7(1), 18–22. https://doi.org/10.1016/0955-0674(95)80040-9

Sun, G., Zhang, M., Chen, H., & Hochstrasser, M. (2022). The CinB Nuclease from wNo Wolbachia Is Sufficient for Induction of Cytoplasmic Incompatibility in Drosophila. mBio, e0317721. https://doi.org/10.1128/mbio.03177-21

Tang, X., Cao, J., Zhang, L., Huang, Y., Zhang, Q., & Rong, Y. S. (2017). Maternal Haploid, a Metalloprotease Enriched at the Largest Satellite Repeat and Essential for Genome Integrity in Drosophila Embryos. Genetics, 206(4), 1829–1839. https://doi.org/10.1534/genetics.117.200949

Teixeira, F. K., Okuniewska, M., Malone, C. D., Coux, R. X., Rio, D. C., & Lehmann, R. (2017). piRNA-mediated regulation of transposon alternative splicing in the soma and germ line. Nature, 552(7684), 268–272. https://doi.org/10.1038/nature25018

Titen, S. W., & Golic, K. G. (2008). Telomere loss provokes multiple pathways to apoptosis and produces genomic instability in Drosophila melanogaster. Genetics, 180(4), 1821–1832. https://doi.org/10.1534/genetics.108.093625

Tram, U., Fredrick, K., Werren, J. H., & Sullivan, W. (2006). Paternal chromosome segregation during the first mitotic division determines Wolbachia-induced cytoplasmic incompatibility phenotype. J Cell Sci, 119(Pt 17), 3655–3663. https://doi.org/10.1242/jcs.03095

Tram, U., & Sullivan, W. (2002). Role of delayed nuclear envelope breakdown and mitosis in Wolbachia-induced cytoplasmic incompatibility. Science, 296(5570), 1124–1126. https://doi.org/10.1126/science.1070536

Turelli, M., & Hoffmann, A. A. (1991). Rapid spread of an inherited incompatibility factor in California Drosophila. Nature, 353(6343), 440–442. https://doi.org/10.1038/353440a0

Wang, H., Xiao, Y., Chen, X., Zhang, M., Sun, G., Wang, F., Wang, L., Zhang, H., Zhang, X., Yang, X., Li, W., Wei, Y., Yao, D., Zhang, B., Li, J., Cui, W., Wang, F., Chen, C., Shen, W., Su, D., Bai, F., Huang, J., Ye, S., Zhang, L., Ji, X., Wang, W., Wang, Z., Hochstrasser, M., & Yang, H. (2022). Crystal Structures of Wolbachia CidA and CidB Reveal Determinants of Bacteria-induced Cytoplasmic Incompatibility and Rescue. Nat Commun, 13(1), 1608. https://doi.org/10.1038/s41467-022-29273-w

Weinert, L. A., Araujo-Jnr, E. V., Ahmed, M. Z., & Welch, J. J. (2015). The incidence of bacterial endosymbionts in terrestrial arthropods. Proc Biol Sci, 282(1807), 20150249. https://doi.org/10.1098/rspb.2015.0249

Werren, J. H., Baldo, L., & Clark, M. E. (2008). Wolbachia: master manipulators of invertebrate biology. Nat Rev Microbiol, 6(10), 741–751. https://doi.org/10.1038/nrmicro1969

Wickham, H. (2016). ggplot2: Elegant Graphics for Data Analysis. In Springer-Verlag. https://ggplot2/tidyverse.org

Wright, J. D., & Barr, A. R. (1981). Wolbachia and the Normal and Incompatible Eggs of Aedes Polynesiensis (Diptera, Culicidae). Journal of Invertebrate Pathology, 38(3), 409–418. https://doi.org/Doi 10.1016/0022-2011(81)90109-9

Wu, X., Lindsey, A. R. I., Chatterjee, P., Werren, J. H., Stouthamer, R., & Yi, S. V. (2020). Distinct epigenomic and transcriptomic modifications associated with Wolbachia-mediated asexuality. PLoS Pathog, 16(3), e1008397. https://doi.org/10.1371/journal.ppat.1008397

Ye, Y. H., Woolfit, M., Huttley, G. A., Rances, E., Caragata, E. P., Popovici, J., O’Neill, S. L., & McGraw, E. A. (2013). Infection with a Virulent Strain of Wolbachia Disrupts Genome Wide-Patterns of Cytosine Methylation in the Mosquito Aedes aegypti. PLoS One, 8(6), e66482. https://doi.org/10.1371/journal.pone.0066482

Yen, J. H., & Barr, A. R. (1971). New hypothesis of the cause of cytoplasmic incompatibility in Culex pipiens L. Nature, 232(5313), 657–658. https://doi.org/10.1038/232657a0

Yuan, K., Seller, C. A., Shermoen, A. W., & O’Farrell, P. H. (2016). Timing the Drosophila Mid-Blastula Transition: A Cell Cycle-Centered View. Trends Genet, 32(8), 496–507. https://doi.org/10.1016/j.tig.2016.05.006

Zheng, X., Zhang, D., Li, Y., Yang, C., Wu, Y., Liang, X., Liang, Y., Pan, X., Hu, L., Sun, Q., Wang, X., Wei, Y., Zhu, J., Qian, W., Yan, Z., Parker, A. G., Gilles, J. R. L., Bourtzis, K., Bouyer, J., Tang, M., Zheng, B., Yu, J., Liu, J., Zhuang, J., Hu, Z., Zhang, M., Gong, J. T., Hong, X. Y., Zhang, Z., Lin, L., Liu, Q., Hu, Z., Wu, Z., Baton, L. A., Hoffmann, A. A., & Xi, Z. (2019). Incompatible and sterile insect techniques combined eliminate mosquitoes. Nature, 572(7767), 56–61. https://doi.org/10.1038/s41586-019-1407-9

